# Deciphering the Agonist Binding Mechanism to the Adenosine A1 Receptor

**DOI:** 10.1101/2020.10.22.350827

**Authors:** Giuseppe Deganutti, Kerry Barkan, Barbara Preti, Michele Leuenberger, Mark Wall, Bruno Frenguelli, Martin Lochner, Graham Ladds, Christopher A Reynolds

## Abstract

Despite being amongst the most characterized G protein-coupled receptors (GPCRs), adenosine receptors (ARs) have always been a difficult target in drug design. To date, no agonist other than the natural effector and the diagnostic regadenoson has been approved for human use. Recently, the structure of the adenosine A1 receptor (A_1_R) was determined in the active, G_i_ protein complexed state; this has important repercussions for structure-based drug design. Here, we employed supervised molecular dynamics simulations and mutagenesis experiments to extend the structural knowledge of the binding of selective agonists to A_1_R. Our results identify new residues involved in the association and dissociation pathway, suggest the binding mode of N6-cyclopentyladenosine (CPA) related ligands, and highlight the dramatic effect that chemical modifications can have on the overall binding mechanism.

## 1 Introduction

The dephosphorylation of adenosine triphosphate (ATP), diphosphate (ADP), and monophosphate (AMP) produces adenosine^1^, a nucleoside present in extracellular concentrations of 20-300 nM under physiological conditions^2^. Adenosine acts ubiquitously on four different G protein-coupled receptors (GPCRs), the adenosine receptor 1 (A_1_R), 2A (A_2A_R), 2B (A_2B_R) and 3 (A_3_R) contributing to the broad range of purinergic signalling^3,4^. The main effect mediated by adenosine comprises of the inhibition or stimulation of adenylate cyclase, the activation of phospholipase C, intracellular Ca^2+^ regulation, and the mitogen-activated protein kinases (MAPK) pathways^5^.

Agonist-activated A_1_R couples to inhibitory G proteins (G_i/o_) to trigger numerous physiological effects. For example, it produces negative chronotropic and inotropic effects in the heart^6^, reduces the neuronal firing rate by blocking neurotransmitter release in the central nervous system (CNS), and inhibits lipolysis and renin release^7^. Therapeutic approaches based on stimulating the A_1_R could pave the way for hitherto unavailable tools to treat the central nervous system (CNS) and cardiovascular diseases^5,8^. From this standpoint, the structure-based drug design (SBDD) of novel agonists can exploit the recent cryo-EM structure of A_1_R in complex with both G_i2_ and adenosine^9^. Like all the other GPCRs, A_1_R presents a transmembrane domain (TMD) formed by seven α-helixes spanning the cytosolic membrane and shaping the orthosteric and the intracellular G_i_ protein binding sites (Figure S1a). Three intracellular loops (ICL1-3) and three extracellular loops (ECL1-3) interconnect the TM helices. ECL2, the longest A_1_R loop, orients almost perpendicularly to the plane of the membrane both in the active^9^ and inactive^10,11^ A_1_R states, in contrast to ECL2 of A_2A_R, which is almost parallel to the membrane^10^. ECL2 is important for A_1_R ligands^12^; it has been implicated in the intermediate states that anticipate the orthosteric complex, therefore acting as a selectivity filter. Moreover, positive allosteric modulators can to bind to it^13,14^.

The key orthosteric interactions between adenosine and A_1_R (Figure S1b) are hydrogen bonds with residues N254^6.55^, S277^7.42^, H278^7.43^, and a π-π interaction that involves F171^ECL2^. AR agonists bearing small N6-cycloalkyl groups, like the N6-cyclopentyladenosine (CPA, Figure 1), display A_1_R selectivity^15^. It is proposed that the ligand selectivity for A_1_R is driven by a small hydrophobic pocket underneath ECL3, due to the presence of T270^7.35^ in place of M270^7.35^ (A_2A_)^16,9,10^.

**Figure 1.**
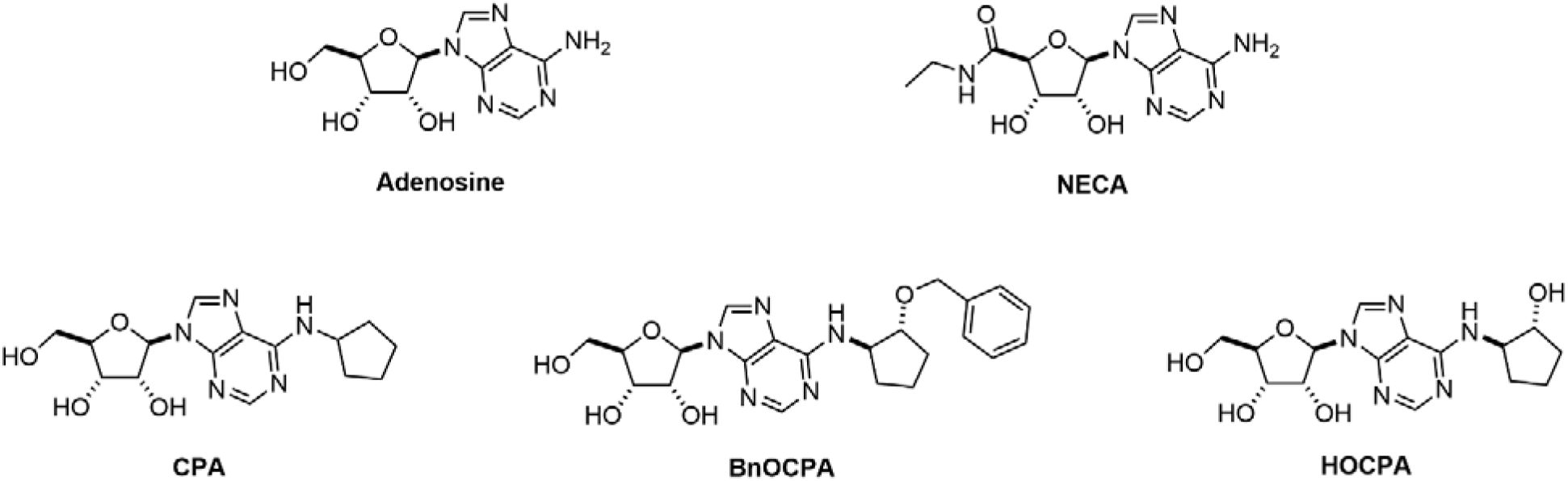
A_1_R agonists considered. Adenosine is the endogenous effector; NECA is a nonselective exogenous ARs agonist; CPA represent a prototypical A_1_R selective agonist; HOCPA and BnOCPA are A_1_R selective analogues of CPA.

Here, we extensively studied the A_1_R recognition of adenosine, 5’-N-carboxamidoadenosine (NECA), CPA, and the recently characterized agonists HOCPA and BnOCPA^17^ (Figure 1). Binding and unbinding pathways were simulated by means of supervised molecular dynamics (SuMD)^18,19^ and the outcomes tested in mutagenesis experiments to identify A_1_R residues involved along the route towards and from the orthosteric site. We propose the binding conformation of N6-cyclopenthyl agonists, and how chemical modifications could impact the binding mechanisms of these selective A_1_R agonists. Our results can be framed within the dynamic nature of ligand-receptor complex formations and highlight the dramatic effect on drug binding kinetics triggered by chemical substitutions.

## 2 Methods

### 2.1 Experimental Methods

#### 2.1.1 Compounds

Adenosine, NECA ((2S,3S,4R,5R)-5-(6-aminopurin-9-yl)-N-ethyl-3,4-dihydroxyoxolane-2-carboxamide), CPA, were purchased from Sigma-Aldrich and dissolved in dimethyl-sulphoxide (DMSO). HOCPA and BnOCPA was synthesised as described in Knight et al., 2016 (compounds 6 and 7 respectfully). CA200645, a high affinity AR xanthine amine congener (XAC) derivative containing a polyamide linker connected to the BY630 fluorophore, was purchased from HelloBio (Bristol, UK) and dissolved in DMSO. The concentration of DMSO was maintained to 1.1% for NanoBRET ligand-binding experiments using CA200645.

#### 2.1.2 Generation of mutant A_1_R constructs

The NanoLluc(Nluc)-tagged human A_1_R pcDNA3.1+ construct used to generate stable HEK 293 cell lines was kindly gifted to us by Stephen Hill and Stephen Briddon (University of Nottingham). Mutations within the A_1_R were made using the QuikChange Lightening Site-Directed Mutagenesis Kit (Agilent Technologies) in accordance with the manufacturer’s instructions. All oligonucleotides used for mutagenesis were designed using the online Agilent Genomics ‘QuikChange Primer Design’ tool and purchased from Merck. All constructs were confirmed by in-house Sanger sequencing.

#### 2.1.3 Cell culture and transfection’s

HEK 293 cells in a single well of 6-well plate (confluency ≥ 80%) were transfected with 2 μg of DNA using polyethyleneimine (PEI, 1 mg/ml, MW = 25,000 g/mol) (Polysciences Inc) at a DNA:PEI ratio of 1:6 (w/v). Briefly, DNA and PEI were added to separate sterile tubes containing 150 mM sodium chloride (NaCl) (total volume 50 μl), allowed to incubate at room temperature for 5 minutes, mixing together and incubating for a further 10 minutes prior to adding the combined mix dropwise to the cells. 48 hours post-transfection, stable Nluc-A_1_R expressing HEK 293 cells were selected using 600 μg/mL Geneticin (Thermo Fisher Scientific) whereby the media was changed every two days. HEK 293 cell lines were routinely cultured in DMEM/F-12 GlutaMAX (Thermo Fisher Scientific) supplemented with 10% FBS (F9665, Sigma-Aldrich).

#### 2.1.4 Analysis of A_1_R cell-surface expression using flow cytometry

HEK 293 cells and WT or mutant A_1_R expressing HEK 293 cells were harvested using a non-enzymatic cell dissociation solution (Sigma-Aldrich) and washed with PBS prior to counting. 0.5 × 10^6^ cells were washed three times in flow buffer (PBS supplemented with 1% BSA and 0.03% sodium azide) before re-suspending in 50 μl flow buffer containing anti-Nluc polyclonal primary antibody raised in rabbit (Kindly gifted by Promega) at 1:100 dilution and incubated at room temperature for 1 hour. All samples were washed three times with flow buffer and re-suspended in 50 μl flow buffer containing Allophycocyanin (APC)-conjugated anti-Rabbit IgG (Thermo Fisher Scientific, 31984) at 1:150 dilution and incubated for 1 hour at room temperature in the dark. The cells received a final three washes and were re-suspended in 300 μl flow buffer.

Analysis was conducted using a BD AccuriTM C6 Plus Flow Cytometer which is equipped with a blue (488 nm) and red (640 nm) laser, two light scatter detectors (FSC and SCC) and four fluorescence detectors (FL1 Em. λ 530/30 nm, FL2 Em. λ 585/40 nm, FL3 Em. λ 570 and FL4 Em. λ 675/25 nm). FL4 optical filters were chosen APC (Ex. λ 633 nm and Em. λ 660 nn). Unstained cells and HEK 293 cells without A_1_R expression were used as controls for autofluorescence and unspecific antibody binding, respectively. All data is collected by the flow cytometer and analysis can be conducted at any time using the BD AccuriTM C6 software. The images for figures were made in FlowJo^®^ (V7.6.5), an analysis platform for single-cell flow cytometry analysis.

#### 2.1.5 BRET assays for binding

Saturation binding at WT and mutant A_1_R for the determination of CA200645 K_D_ was conducted using NanoBRET. The Nluc acts as the BRET donor (luciferase oxidizing its substrate, furimazine) and CA200645 acted as the fluorescent acceptor. Here, HEK 293 cells stably expressing WT or mutant Nluc-A_1_R 24 hours post plating (10,000 cells/well of a 96-well plate) where pre-treated with 0.1 μM furimazine for 5 minutes prior to stimulated with CA200645 at a range of concentrations (0 to 300 nM). Following a 30 minute incubation period at room temperature, filtered light emission at 450 nm and > 610 nm (640-685 nm band pass filter) was measured using a Mithras LB 940 and the raw BRET ratio calculated (610 nm/450 nm). Non-specific binding was determined by saturating concentrations of DPCPX (1 μM) and subtracted from the BRET ratio. BRET ratio data was then fit to the ‘One site – Specific binding’ model built into Prism.

As described previously (Barkan et al. 2019), NanoBRET competition binding assays were conducted to determine the affinity (pKi) of various A_1_R compounds. Briefly, following a 5 minute pre-incubation with 0.1 μM furimazine, cells were co-stimulated with CA200645 (used at 25 nM, as previously reported (Stoddart et al. 2015; Barkan et al. 2019)) and increasing concentration of unlabeled ligand and emission at 450 nm and > 610 nm immediately measured. The BRET ratio at 10 minutes post stimulation was fitted with the ‘one-site – Ki model’ derived from the Cheng and Prusoff correction (Chen and Prusoff, 1973), built into Prism to determine affinity (pKi) values for all unlabeled agonists at the A_1_Rs. The determined Kd of CA200645 at the mutant A_1_R was taken into account during analysis. Nonspecific binding was determined using a high concentration of unlabelled antagonist, at 1 μM DPCPX.

#### 2.1.6 Data analysis

All experiments were conducted in duplicate (technical replicates) to ensure the reliability of single values. Statistical analysis, performed using Prism 8.0, was undertaken for experiments where the group size was at least n = 3 and these independent values used to calculate statistical significance (*, *p*<*0.05*; **, *p*<*0.01*; ***, *p*<*0.001*; ****, *p*<*0.0001*) using a one-way ANOVA with a Dunnett’s post-test for multiple comparisons.

### 2.2 Computational Methods

#### 2.2.1 Biological targets and ligands force field parameters

All ten systems (Table 1) were prepared for molecular dynamics (MD) simulations using the CHARMM36^20,21^/CGenFF 3.0.1^22–24^ force field combination. Initial ligand force filed topology and parameter files were obtained from the ParamChem webserver^22^. Adenosine and NECA are already well-parameterized in the CGenFF force filed. Optimized parameters for HOCPA and BnOCPA from our previous work^25^ were used and transferred to CPA.

**Table 1.** The ten systems simulated employing SuMD and SuMD path sampling.

| **Ligand** | **Transition** | **# SuMD replicas** | **Total SuMD path sampling** |
| --- | --- | --- | --- |
| **Adenosine** | Binding | 9 | 5.18 μs |
| | Unbinding | 5 | 3.09 μs |
| **NECA** | Binding | 6 | 1.86 μs |
| | Unbinding | 6 | 7.10 μs |
| **CPA** | Binding | 9 | 4.34 μs |
| | Unbinding | 10 | 0.97 μs |
| **HOCPA** | Binding | 8 | 4.06 μs |
| | Unbinding | 6 | 2.58 μs |
| **BnOCPA** | Binding | 7 | 5.82 μs |
| | Unbinding | 6 | 4.92 μs |

#### 2.2.2 Protein preparation

The active A_1_R structure (Table 1) was retrieved from the Protein Data Bank^26^ (PDB code 6D9H^9^). For SuMD binding simulations, the agonists were placed at least 30 Å from the binding site in five different systems. For SuMD unbinding, the experimental coordinates (PDB 6D9H) were used to simulate the adenosine, while representative frames (ligand orthosteric conformations close to the experimental bound adenosine on A_1_R) from the SuMD binding were used to start the SuMD unbinding of NECA, CPA, HOCPA and BnOCPA. The A_1_R intracellular loop 3 (ICL3) was modelled using Modeller 9.19^27^. For all the 10 systems (Table 1), hydrogen atoms were added by means of the pdb2pqr^28^ and propka^29^ software (considering a simulated pH of 7.0); the protonation of titratable side chains was checked by visual inspection. The resulting receptor was inserted in a square 90 Å × 90 Å 1-palmitoyl-2-oleyl-sn-glycerol-3-phosphocholine (POPC) bilayer (previously built by using the VMD Membrane Builder plugin 1.1, Membrane Plugin, Version 1.1 at: http://www.ks.uiuc.edu/Research/vmd/plugins/membrane/), through an insertion method^30^. The receptor orientation was obtained by superposing the coordinates on the corresponding structure retrieved from the OPM database^31^. Lipids overlapping the receptor transmembrane helical bundle were removed and TIP3P water molecules^32^ were added to the simulation box by means of the VMD Solvate plugin 1.5 (Solvate Plugin, Version 1.5. at <http://www.ks.uiuc.edu/Research/vmd/plugins/solvate/). Finally, overall charge neutrality was reached by adding Na^+^/Cl^−^ counter ions up to the final concentration of 0.150 M), using the VMD Autoionize plugin 1.3 (Autoionize Plugin, Version 1.3. at <http://www.ks.uiuc.edu/Research/vmd/plugins/autoionize/).

#### 2.2.3 Systems equilibration and general MD settings

The MD engine ACEMD^33^ was employed for both the equilibration and productive simulations. The equilibration of the membrane systems was achieved in isothermal-isobaric conditions (NPT) using the Berendsen barostat^34^ (target pressure 1 atm) and the Langevin thermostat^35^ (target temperature 300 K) with low damping of 1 ps^−1^. A four-stage procedure was performed (integration time step of 2 fs): first, clashes between protein and lipid atoms were reduced through 2000 conjugate-gradient minimization steps, then a 2 ns long MD simulation was run with a positional constraint of 1 kcal mol^−1^ Å^−2^ on protein and lipid phosphorus atoms. During the second stage, 20 ns of MD simulation were performed constraining only the protein atoms, while in the last equilibration stage, positional constraints were applied only to the protein backbone alpha carbons, for a further 20 ns. Globular protein equilibration was achieved in two steps: after 500 cycles of conjugate-gradient minimization, the system was simulated for 5 ns, employing an integration time step of 2 fs, in the isothermal-isobaric conditions (NPT).

Productive trajectories (Table S4) were computed with an integration time step of 4 fs in the canonical ensemble (NVT). The target temperature was set at 300 K, using a thermostat damping of 0.1 ps^−1^; the M-SHAKE algorithm^36,37^ was employed to constrain the bond lengths involving hydrogen atoms. The cut-off distance for electrostatic interactions was set at 9 Å, with a switching function applied beyond 7.5 Å. Long-range Coulomb interactions were handled using the particle mesh Ewald summation method (PME)^38^ by setting the mesh spacing to 1.0 Å.

#### 2.2.4 The supervised MD (SuMD) protocol

The supervised molecular dynamics (SuMD) is an adaptive sampling method^39^ for speeding up the simulation of binding^18,40^ and unbinding processes^19,41^. In the simplest SuMD implementation, sampling is gained without the introduction of any energetic bias, by applying a tabu–like algorithm to monitor the distance between the centers of mass (or the geometrical centers) of the ligand and the predicted binding site or the receptor. However, the supervision of a second metric of the system can be considered^41^. A series of short unbiased MD simulations are performed, and after each simulation, the distances (collected at regular time intervals) are fitted to a linear function. If the resulting slope is negative (for binding) or positive (for unbinding) the next simulation step starts from the last set of coordinates and velocities produced, otherwise, the simulation is restarted by randomly assigning the atomic velocities.

#### 2.2.5 Settings for SuMD binding to the A_1_R

To simulate the agonists’ binding to the A_1_R (Table 1, Video S1-S5) the distance between the centroid of the ligand and the centroid of the orthosteric residues N254^6.55^, F171^ECL2^, T277^7.42^, and H278^7.43^ was supervised during 500 ns long time windows until it reached a value less than 4 Å. Then?

#### 2.2.6 Settings for SuMD unbinding from the A_1_R

For the SuMD unbinding (Table 1, Video S1-S4), differently from the SuMD binding algorithm, the length (Δt) of the short simulations increases along the dissociation pathway, according to the formula:

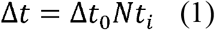

Δt_0_ is the duration of the very first MD time window and Nt_i_ is a factor that is picked from three user-defined values (Nt_1_, Nt_2_, and Nt_3_), according to the last ligand-protein distance detected. At the end of each MD run, the ligand-protein distance (r_L_) is compared to three distance threshold values (D_1_, D_2_ and D_3_, also defined by the user), allowing a decision on the value of Nt_i_ factor according to the following conditions:

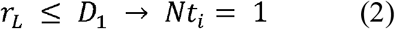

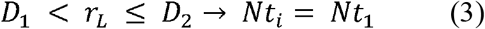

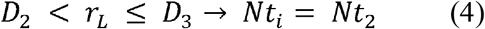

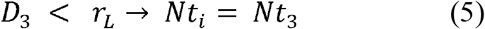

Values of 3 Å, 5 Å and 8 Å were used for D_1_, D_2_ and D_3_ respectively, while Nt_1_, Nt_2_, and Nt_3_ were set to 2, 4, and 8 (SuMD time windows of 100 ps, 200 ps, 400 ps and 800 ps).

The adenosine SuMD binding and unbinding trajectories were the same as our previous work^19^.

#### 2.2.7 SuMD path sampling protocol

Further MD sampling (SuMD path sampling, Table 1) was performed using the outputs from each SuMD replica (Video S1-S4) for both binding and unbinding. Each trajectory was aligned on the protein alpha carbon atoms and the frames were clustered according to the ligand root mean square deviation (RMSD) to the starting positions (bin of 1 Å). A frame from each group was randomly extracted and used as a starting point for 20 ns long classic MD simulations.

#### 2.2.8 Analysis of the MD trajectories

Only the MD trajectories from the SuMD path sampling were analyzed. Interatomic contacts and root mean square deviations (RMSD) were computed using VMD^42^. Contacts were considered productive if the distance between two atoms was less than 3.5 Å. Ligand-protein hydrogen bonds were detected using the GetContacts scripts tool (https://getcontacts.github.io), setting a hydrogen bond donor-acceptor distance of 3.3 Å and an angle value of 150° as geometrical cut-offs. Contacts and hydrogen bond persistency are quantified as the percentage of frames (considering all the frames obtained by merging the different replicas) in which protein residues formed contacts or hydrogen bonds with the ligand. The computation takes into account direct and water-mediated interactions.

Distances between atoms were computed using PLUMED 2.3^43^. The molecular mechanics generalized Born surface area (MM/GBSA) energy was computed with the MMPBSA.py^44^ script (AmberTools17 suite at http://ambermd.org/), after transforming the CHARMM psf topology files to an Amber prmtop format using ParmEd (documentation. at <http://parmed.github.io/ParmEd/html/index.html). We preferred the MM/GBSA approach over the molecular Poisson-Boltzmann surface area (MM/PBSA) because binding and unbinding paths are not compatible with a grid method^45^.

### 2.3 Numbering system

Throughout the manuscript, the Ballesteros-Weinstein residues numbering system for the GPCRs^46^ is adopted as superscript.

## 3 Results and Discussion

### 3.1 N6-cyclopentyl agonists bind to A_1_R with similar fashion

ARs ligands bearing an alkyl ring in N6 (on the adenine scaffold) or C8 (xanthine scaffold) position display increased affinity for A_1_R over A_2A_R^10^. The reason for this is attributed to T270^7.35^ (A_1_R) in place of M270^7.35^ (A_2A_R), which shapes a hydrophobic sub-pocket underneath ECL3 that partially accommodates the lipophilic substituent. While the inactive structures of A_1_R and A_2A_R in complex with the xanthine antagonist PSB36^10^ have univocally shown this structural aspect of the selectivity, no confirmation from crystallography or cryo-EM studies is yet available for ARs agonists.

Besides reproducing the adenosine cryo-EM binding conformation into A_1_R (Figure S2, Video S1)^9^, our simulations sampled the orthosteric binding mode of NECA observed on A_2A_R (Figure S2, Video S2)^47^. In light of this reliability, we propose the likely binding mode of CPA, HOCPA, and BnOCPA (Figure 2, Videos S3-S4). As expected, the adenine ring forms a bidentate hydrogen bond with N254^6.55^ and a π- π stacking with F171^ECL2^, while the N6-cyclopentyl ring inserts in the hydrophobic pocket under ECL3, interacting with T270^7.35^ and L253^6.54^ (Figure 2). Mutagenesis experiments confirmed the importance of L253^6.53^ (Table 2) for the affinity of the agonists. BnOCPA is proposed to bind to A_1_R with same features of the smaller ligands CPA and HOCPA (Figure 2b, Video S4). However, during the simulations the oxybenzyl group showed^25^ high flexibility and explored three different orientations (Figure 2b). In two of these conformations, BnOCPA interacted with A_1_R residue L258^6.59^ (mode B, Figure 2b) and Y271^7.36^ (mode C, Figure 2b). Binding assays with mutants L258^6.59^A, F258^6.59^A, F258^6.59^T and Y271^7.36^A (Table 2) confirmed that these two residues are likely involved in the orthosteric complex with BnOCPA. However, a role during agonist association and dissociation events cannot be ruled out, as also the affinity for CPA and HOCPA (which are not involved in contacts with L258^6.59^ and Y271^7.36^ in the bound state, Figure 2a) was affected (Table 2).

**Figure 2.**
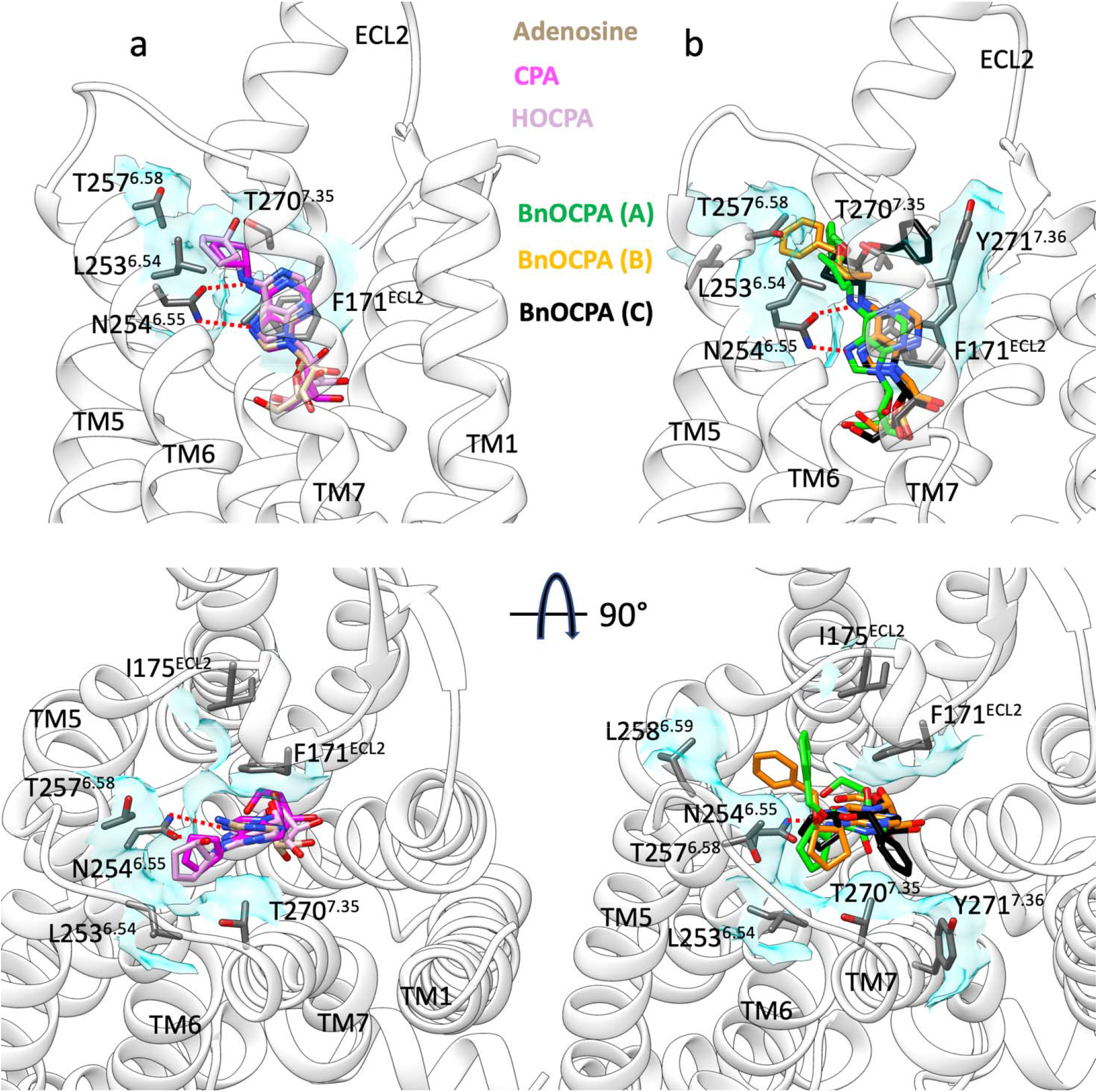
Binding modes of the agonists according to SuMD path sampling simulations. **a**) CPA (magenta) and HOCPA (pink) engage A_1_R with the same orientation as adenosine (tan stick); the N6-cyclopentyl group interacts with L253^6.54^ and T257^6.58^. **b**) BnOCPA orients the oxybenzyl group in three different orientations. Hydrogen bonds with N254^ECL2^ are shown as dashed lines, while hydrophobic contacts are depicted as cyan transparent surfaces.

**Table 2.**
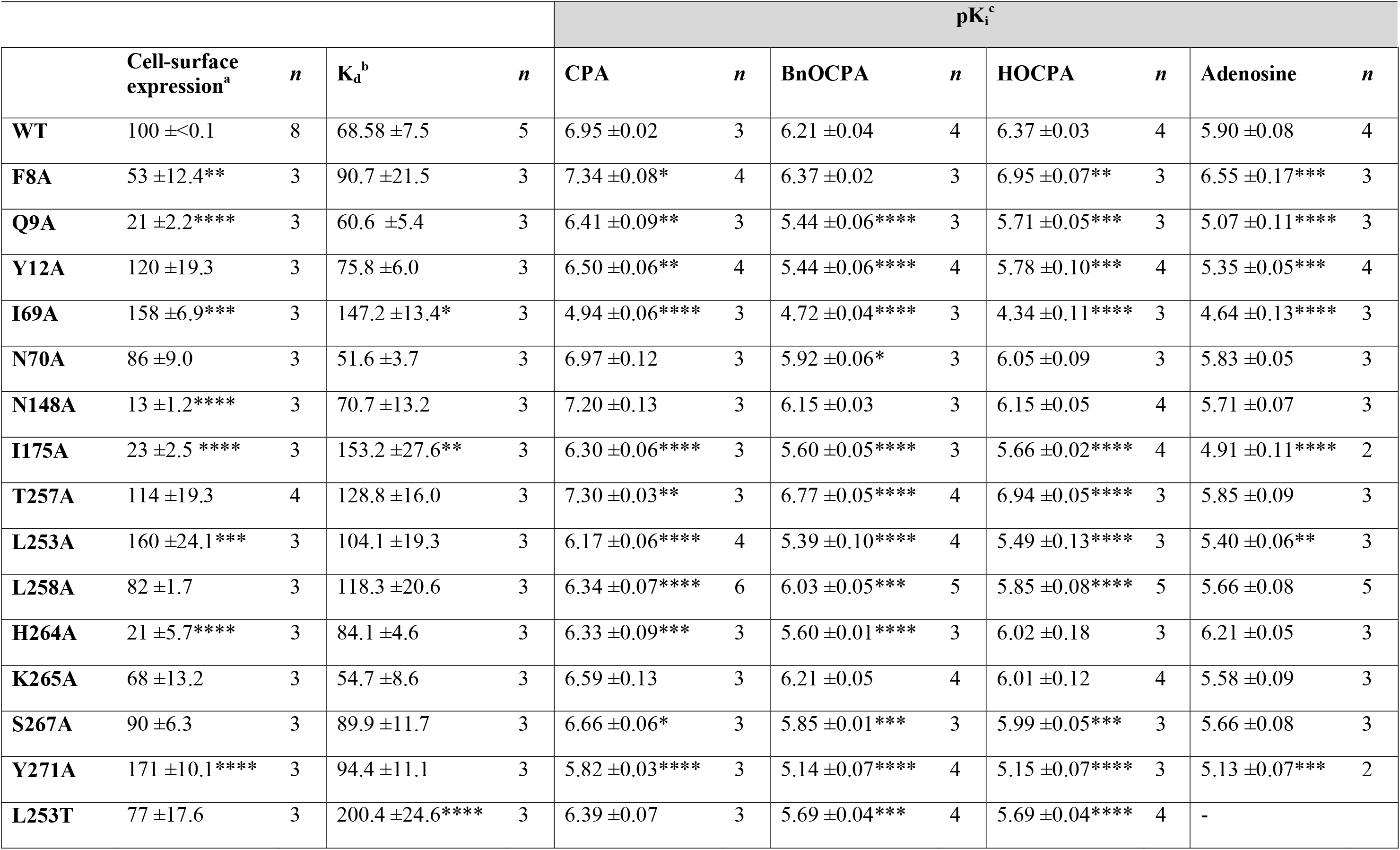
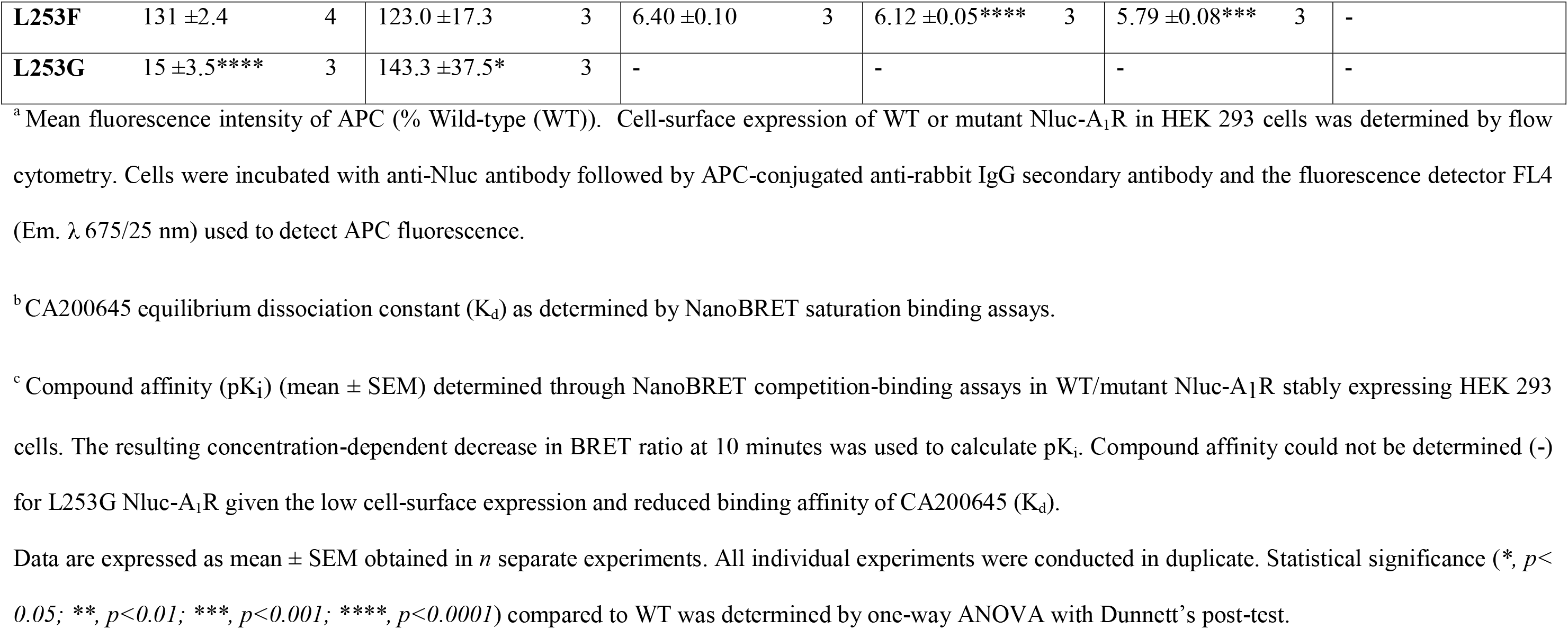
NanoBRET saturation- and competition-binding assays in WT and mutant Nluc-A_1_R. CA200645 equilibrium dissociation constant (K_d_) and compound affinity (pK_i_) at WT and mutant Nluc-A_1_R, as determined by NanoBRET saturation and competition ligand-binding assays, respectively.

|  |  |  |  | pK<sub>i</sub><sup>c</sup> |  |  |  |  |  |  |  |  |  |  |  |  |  |
| --- | --- | --- | --- | --- | --- | --- | --- | --- | --- | --- | --- | --- | --- | --- | --- | --- | --- |
|  | Cell-surface expression<sup>a</sup> | *n* | K<sub>d</sub><sup>b</sup> |  | *n* | CPA |  | *n* | BnOCPA |  | *n* | HOCPA |  | *n* | Adenosine |  | *n* |
| WT | 100 ±<0.1 | 8 | 68.58 ±7.5 |  | 5 | 6.95 ±0.02 |  | 3 | 6.21 ±0.04 |  | 4 | 6.37 ±0.03 |  | 4 | 5.90 ±0.08 |  | 4 |
| F8A | 53 ±12.4** | 3 | 90.7 ±21.5 |  | 3 | 7.34 ±0.08* |  | 4 | 6.37 ±0.02 |  | 3 | 6.95 ±0.07** |  | 3 | 6.55 ±0.17*** |  | 3 |
| Q9A | 21 ±2.2**** | 3 | 60.6 ±5.4 |  | 3 | 6.41 ±0.09** |  | 3 | 5.44 ±0.06**** |  | 3 | 5.71 ±0.05*** |  | 3 | 5.07 ±0.11**** |  | 3 |
| Y12A | 120 ±19.3 | 3 | 75.8 ±6.0 |  | 3 | 6.50 ±0.06** |  | 4 | 5.44 ±0.06**** |  | 4 | 5.78 ±0.10*** |  | 4 | 5.35 ±0.05*** |  | 4 |
| I69A | 158 ±6.9*** | 3 | 147.2 ±13.4* |  | 3 | 4.94 ±0.06**** |  | 3 | 4.72 ±0.04**** |  | 3 | 4.34 ±0.11**** |  | 3 | 4.64 ±0.13**** |  | 3 |
| N70A | 86 ±9.0 | 3 | 51.6 ±3.7 |  | 3 | 6.97 ±0.12 |  | 3 | 5.92 ±0.06* |  | 3 | 6.05 ±0.09 |  | 3 | 5.83 ±0.05 |  | 3 |
| N148A | 13 ±1.2**** | 3 | 70.7 ±13.2 |  | 3 | 7.20 ±0.13 |  | 3 | 6.15 ±0.03 |  | 3 | 6.15 ±0.05 |  | 4 | 5.71 ±0.07 |  | 3 |
| I175A | 23 ±2.5 **** | 3 | 153.2 ±27.6** |  | 3 | 6.30 ±0.06**** |  | 3 | 5.60 ±0.05**** |  | 3 | 5.66 ±0.02**** |  | 4 | 4.91 ±0.11**** |  | 2 |
| T257A | 114 ±19.3 | 4 | 128.8 ±16.0 |  | 3 | 7.30 ±0.03** |  | 3 | 6.77 ±0.05**** |  | 4 | 6.94 ±0.05**** |  | 3 | 5.85 ±0.09 |  | 3 |
| L253A | 160 ±24.1*** | 3 | 104.1 ±19.3 |  | 3 | 6.17 ±0.06**** |  | 4 | 5.39 ±0.10**** |  | 4 | 5.49 ±0.13**** |  | 3 | 5.40 ±0.06** |  | 3 |
| L258A | 82 ±1.7 | 3 | 118.3 ±20.6 |  | 3 | 6.34 ±0.07**** |  | 6 | 6.03 ±0.05*** |  | 5 | 5.85 ±0.08**** |  | 5 | 5.66 ±0.08 |  | 5 |
| H264A | 21 ±5.7**** | 3 | 84.1 ±4.6 |  | 3 | 6.33 ±0.09*** |  | 3 | 5.60 ±0.01**** |  | 3 | 6.02 ±0.18 |  | 3 | 6.21 ±0.05 |  | 3 |
| K265A | 68 ±13.2 | 3 | 54.7 ±8.6 |  | 3 | 6.59 ±0.13 |  | 3 | 6.21 ±0.05 |  | 4 | 6.01 ±0.12 |  | 4 | 5.58 ±0.09 |  | 3 |
| S267A | 90 ±6.3 | 3 | 89.9 ±11.7 |  | 3 | 6.66 ±0.06* |  | 3 | 5.85 ±0.01*** |  | 3 | 5.99 ±0.05*** |  | 3 | 5.66 ±0.08 |  | 3 |
| Y271A | 171 ±10.1**** | 3 | 94.4 ±11.1 |  | 3 | 5.82 ±0.03**** |  | 3 | 5.14 ±0.07**** |  | 4 | 5.15 ±0.07**** |  | 3 | 5.13 ±0.07*** |  | 2 |
| L253T | 77 ±17.6 | 3 | 200.4 ±24.6**** |  | 3 | 6.39 ±0.07 |  | 3 | 5.69 ±0.04*** |  | 4 | 5.69 ±0.04**** |  | 4 | - |  |  |

|  |  |  |  |  |  |  |  |  |  |  |  |
| --- | --- | --- | --- | --- | --- | --- | --- | --- | --- | --- | --- |
| **L253F** | 131 ±2.4 | 4 | 123.0 ±17.3 | 3 | 6.40 ±0.10 | 3 | 6.12 ±0.05**** | 3 | 5.79 ±0.08*** | 3 | - |
| **L253G** | 15 ±3.5**** | 3 | 143.3 ±37.5* | 3 | - |  | - |  | - |  | - |

<sup>a</sup> Mean fluorescence intensity of APC (% Wild-type (WT)). Cell-surface expression of WT or mutant Nluc-A<sub>1</sub>R in HEK 293 cells was determined by flow cytometry. Cells were incubated with anti-Nluc antibody followed by APC-conjugated anti-rabbit IgG secondary antibody and the fluorescence detector FL4 (Em. $\lambda$ 675/25 nm) used to detect APC fluorescence.
<sup>b</sup> CA200645 equilibrium dissociation constant ( $K_d$ ) as determined by NanoBRET saturation binding assays.
<sup>c</sup> Compound affinity ( $pK_i$ ) (mean ± SEM) determined through NanoBRET competition-binding assays in WT/mutant Nluc-A<sub>1</sub>R stably expressing HEK 293 cells. The resulting concentration-dependent decrease in BRET ratio at 10 minutes was used to calculate $pK_i$ . Compound affinity could not be determined (-) for L253G Nluc-A<sub>1</sub>R given the low cell-surface expression and reduced binding affinity of CA200645 ( $K_d$ ).
Data are expressed as mean ± SEM obtained in *n* separate experiments. All individual experiments were conducted in duplicate. Statistical significance (\*, $p < 0.05$ ; \*\*, $p < 0.01$ ; \*\*\*, $p < 0.001$ ; \*\*\*\*, $p < 0.0001$ ) compared to WT was determined by one-way ANOVA with Dunnett's post-test.

### 3.2 Simulations suggest the binding and unbinding paths of A_1_R agonists

SuMD and SuMD path sampling delivered detailed insights on the possible transitory metastable states that the agonists experienced along the route to, and from, the orthosteric site (Figure 3, Table S1-S4, Figure S4, Figure S5; Video S1-S4). While the endogenous agonist adenosine reached the orthosteric site with very limited intermediate interactions with ECL2 (Figure 3b, Table S1, Figure S4, Video S1), the other ligands formed contacts with this extracellular vestibule, with CPA and BnOCPA most involved in metastable states (Figure 3d,h, Table S1, Figure S4, Video S3, Video S4). As a general view, the increase in lipophilicity at the N6 position (CPA, HOCPA and BnOCPA, Figure 3d,f,h, Table S1, Figure S4) and 5’ position (NECA, Figure 3, Figure S3a, Table S1, Figure S4) favored intermediate interactions with ECL2. The importance of ECL2 for the binding of NECA, CPA and the antagonist DPCPX to A_1_R has been recently demonstrated^12,48^. Our simulations suggest I175^ECL2^ and E172^ECL2^ as involved in interactions during both the binding and unbinding (Figure 3d, Figure S3, Figure S5), while E170^ECL2^, I167^ECL2^, N159^ECL2^, and W156^ECL2^ engaged NECA during the binding (Figure S3a, Figure S5). L149^ECL2^, on the other hand, was not involved in direct interactions with the ligands, suggesting an important role in stabilizing the overall secondary structure of ECL2 due to its position at the base of the loop helix. Besides the aforementioned residues, further A_1_R side chains were involved along the simulated binding paths for all the agonists considered (Table S1, Table S2 Figure S4).

**Figure 3.**
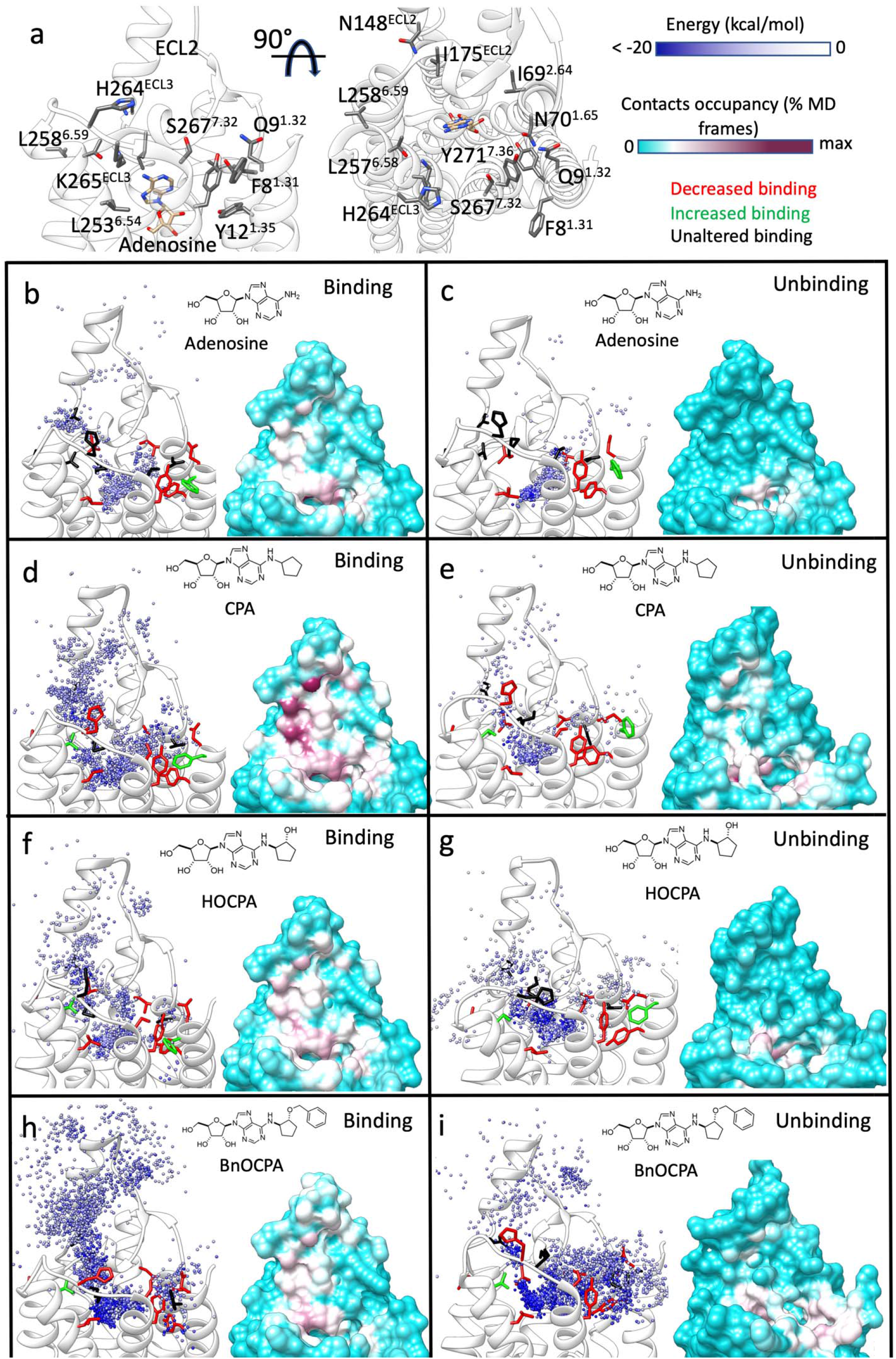
Simulated binding and unbinding paths of the agonists and the relative position of the A_1_R mutants tested. **a**) Two side views of the A_1_R (white transparent ribbon); residues considered for the mutations (Table 2) are shown as grey sticks. The cryo-EM bound adenosine (tan stick) is reported as reference. **b**) to **i**) Left-hand panels, position of the agonist centroid during simulations, colored according to the interaction energy with A_1_R (white ribbon); residues mutated (Table 2) are shown as sticks and colored according to effect on the affinity (red: decreased affinity; green: increased affinity; black: unaltered affinity). Right-hand panels, A_1_R-agonist contacts plotted onto the protein surface and colored according to the contacts occupancy. **b**) Adenosine binding simulations; **c**) adenosine unbinding simulations; **d**) CPA binding simulations; **e**) CPA unbinding simulations; **f**) HOCPA binding simulations; **g**) HOCPA unbinding simulations; **h**) BnOCPA binding simulations; **i**) BnOCPA unbinding simulations.

SuMD unbinding routes (Figure 3c,e,g,I, Video S1-S4) had only limited overlap with the binding routes (Figure 3b,d,f,h, Video S1-S4). The agonists, indeed, established frequent interactions with the top of TM1, TM2, and TM7 (Figure 3c,e,g,i, Figure S3b, Table S2, Table S4, Figure S5). However, in analogy with the binding simulations, CPA, HOCPA, and BnOCPA (all bearing an N6 hydrophobic moiety) showed a major involvement of ECL2 (Table S3, Table S4, Figure S5; Video S3-S4). Many of the A_1_R residues involved in both agonist association (Table S1, Table S2, Figure S4) and dissociation (Table S3, Table S4, Figure S5) are part of the orthosteric site. It is therefore not surprising that mutagenesis experiments already pointed out their importance^12,49–53^.

Interestingly, our unbiased nonequilibrium simulations pointed out several residues, located outside the orthosteric site, involved along the binding and unbinding routes. These residues comprise F8^1.31^, Q9^1.32^, Y12^1.35^, I69^2.64^, N70^2.64^, N148^ECL2^, I175^ECL2^, L253^6.54^, T257^6.58^, L258^6.59^, H264^ECL3^, K265^ECL2^, S267^7.32^, Y271^7.36^ (Figure 3a). Mutagenesis experiments were designed and performed to confirm computational predictions (Figure 3, Table 2, Table S5).

### 3.3 Mutagenesis experiments reveal novel A_1_R residues involved in the binding of agonists

#### 3.3.1 Surface expression of the A_1_R alanine mutants and CA200645 Kd

The cell-surface expression of WT and mutant A_1_R, as determined by mean fluorescence intensity of APC (%WT), was comparable to WT for Y12^1.35^A, N70^2.64^A, T257^6.58^A, L258^6.59^A, K265^ECL2^A, S267^7.32^A, L253^6.54^T and L254F A_1_R. When compared to WT, the cell-surface expression was determined to be significantly reduced for F8A, Q9^1.32^A, N148^ECL2^A, I175^ECL2^A, H264^ECL2^A and L253^6.54^G whereas it was significantly increased for I69^2.65^A, L253^6.54^A and Y271^7.36^A. This enhanced or reduced mutant A_1_R cell-surface expression did not correlate with changes in CA200645 equilibrium dissociation constant (Kd). For example, the Kd for CA200645 determined in L253^6.54^T was significantly increased when compared to WT but showed comparable cell-surface expression. I175 ^ECL2^A and L253^6.54^G also showed an increased Kd but had a reduced cell-surface expression when compared to WT.

Importantly, the determined changes in CPA, BnOCPA, HOCPA or adenosine affinity (pKi) were determined to be independent of changes in cell-surface expression. For example, despite the significantly elevated cell-surface expression of Y271^7.36^A, the determined compound affinity of all four tested compounds was significantly reduced when compared to that determined at WT. This is likely due to the high receptor expression in our system and/or reserve. The only exception was L253^6.54^G whereby compound pKi could not be determined due to the likely combined effect of low cell-surface expression and reduced CA200645 affinity.

#### 3.3.2 General effects of alanine substitutions on agonists affinity

A number of mutations enhanced affinity, while other either decreased affinity or had no overall effect. The mutated residues can be divided into three groups according to their positions relative to the (un)binding paths sampled during the simulations.

#### 3.3.3 TM1, TM2 and TM7 residues

The A_1_R residues located at the top of TM1, TM2 and TM7 (Figure 3a) were generally involved during both the simulated binding (Figure 3b,d,f,h, Figure S4, Figure S5) and unbinding (Figure 3c,e,g,i, Figure S4, Figure S5). Adenosine, CPA, and HOCPA displayed decreased affinity to A_1_R mutants Q9^1.32^A and Y12^1.35^A (Table 2) and enhanced the binding to F8^1.31^A compared to the WT, in agreement with the transitory interactions formed with TM1 during simulations (Figure 3b-g, Table S1-S4, Figure S4, Figure S5). The affinity for the agonists diminished on Q9^1.32^A and Y12^1.35^A, but BnOCPA was not significantly affected by F8^1.31^A. This is apparently in disagreement with simulations, which proposed BnOCPA as the agonist more prone to form metastable states in the proximity of F^1.31^8 during the dissociation from the receptor (Figure 3i). However, the fact that BnOCPA was the ligand most prone to interact with F8 also during the association (Figure 3h) may suggest a degree of compensation between binding and unbinding. That is, the variation of the binding rate can be compensated by an opposite change in the unbinding rate (and vice versa) resulting in unchanged affinity. This could also be the case of N70^2.65^A, which did not alter the affinity of the agonists, except for BnOCPA (Table 2). Binding and unbinding simulations suggested that N70^2.65^ forms numerous interactions with the ligands (Table S1-S4, Figure S4, Figure S5). The adjacent residue I69^2.64^, instead, when mutated to alanine (I69^2.64^A) significantly decreased the affinity of all the agonists (Table 2). Besides being involved in frequent interactions with the agonists (Table S1-S4), I69^2.64^ is packed in hydrophobic contacts with TM3, possibly stabilizing the neighboring part of ECL2.

Moving to TM7, the agonists NECA, CPA, HOCPA, and BnOCPA, but not adenosine, displayed diminished binding to the S267^7.32^A (Table 2, Table S5), while all of them lost affinity to Y271^7.36^A (Table2, Table S5). The bulky Y271^7.36^ side chain, which occupies an important position at the interface between the orthosteric site and the extracellular vestibule, could participate in numerous intermediate interactions along the (un)binding routes.

#### 3.3.4 TM6 residues

The three A_1_R residues mutated on TM6 (L253^6.54^, T257^6.58^, and L258^6.59^, Figure 3a) are part of different protein environments. L253^6.54^ shapes part of the hydrophobic pocket underneath ECL3 partially responsible for A_1_/A_2A_ ligands selectivity (along with L269^7.34^ and T270^7.35^). All the agonists displayed diminished affinity to L253^6.54^A (Table 2), with the N6-substituted agonists CPA, HOCPA, and BnOCPA most affected, in line with the hydrophobic interactions occurring in the bound state (Figure 2). Interestingly, T257^6.58^A increased the affinity of CPA, HOCPA, and BnOCPA. This could be due to an increase in lipophilicity of the protein environment surrounding the cyclopentyl group either in the bound complex or along the unbinding pathways, (Figure 3e,g,i). NECA showed reduced affinity to T257A (Table S5, Figure S3), confirming the importance of hydrophobic N6-substituents for interactions with the top of TM6. L258^6.59^, which is located at the interface with the membrane, is one of the TM5 and TM6 residues shaping a saddle between ECL2 and ECL3, where agonists tended to form metastable interactions along the simulated (un)binding paths (Figure 3b-i, Figure S4, Figure S5). All the agonists (excepted adenosine) displayed a reduced affinity for L258^6.59^A and L258^6.59^T, suggesting that the bulkier ligands may be more prone to interact with this part of A_1_R (BnOCPA was the only ligand proposed to interact at some extent with L258^6.59^ in mode B, Figure 2b). The affinity decrease (Table 2) displayed by HOCPA and BnOCPA for L258^6.59^F likely excludes a destabilization of the neighbor protein structure, as residue F258^6.59^ WT A_2A_R does not change the conformation of the top of TM6.

#### 3.3.5 ECL2 and ECL3 residues

As I175^ECL2^ formed numerous interactions with the agonists during the simulations (Figure 3b-i, Table S1, Table S2, Figure S4, Figure S5), not surprisingly the I175^ECL2^A mutation reduced the affinity of all the ligands (Table 2, Table S5). I175^ECL2^ could contribute to keeping the aromatic side chain of F171^ECL2^ in an appropriate conformation for interacting with the ligands’ adenine ring in the orthosteric complex (Figure S1). On the other hand, N148^ECL2^A did not affect the affinity of the agonists, despite the frequent interactions during the binding simulations (Table 2, Figure 3b,d,f,h, Table S1, Table S2, Figure S4, Figure S5).

While H264^ECL2^A showed diminished affinity (Table 2) for CPA and BnOCPA (which have the most lipophilic N6-group, Figure 1), none of the tested ligands were significantly affected by K265^ECL2^A. As K265^ECL2^ is part of a stable salt bridge with E172^ECL2^, K265^ECL2^A is expected to affect the affinity of the ligands due to the reduced hindrance of the orthosteric site. The subtype A_2A_R bears A265^ECL2^ in place of K265^ECL2^ and the residue involved in the salt bridge with E169^ECL2^ (corresponding to E172^ECL2^ in A_1_R) is H264^ECL2^. The A_2A_R H264^ECL2^A mutant does not display a modified affinity for the antagonists ZM241385, despite the increase in off rate^54^. This could indicate a kinetic compensation due to a faster binding to H264^ECL2^A (A_2A_R) and K265^ECL2^A (A_1_R). A less bulky alanine side chain, indeed, would favor ligand binding and the unbinding to a similar extent.

### 3.4 Towards the definition of the structure-binding/unbinding path relationships of A_1_R agonists

The (un)binding kinetics have a major impact on both pharmacodynamics^55–57^ and pharmacokinetics^58^ of a drug. Even small structural modifications within a congeneric series of ligands modify the kinetics through modified *on* and *off* rates^59^. The reason for this lies in the enormously higher number of different intermediate states that a ligand can experience along (un)binding paths compared to the orthosteric complexes, where it is restrained by intermolecular interactions and steric hindrances. Protein-ligand recognition events have very complex energy landscapes^60–63^ and small changes in either the structure of the ligand or the protein can alter the nature and the position of the transition states along the (un)binding routes^64^. To investigate this aspect on the A_1_R agonists, we compared the A_1_R interaction patterns between adenosine, CPA or BnOCPA (Figure 4) to understand how the introduction of the N-6 cyclopentyl group on the adenosine scaffold and the introduction of the oxybenzyl group on CPA (BnOCPA) could alter the overall (un)binding mechanisms.

**Figure 4.**
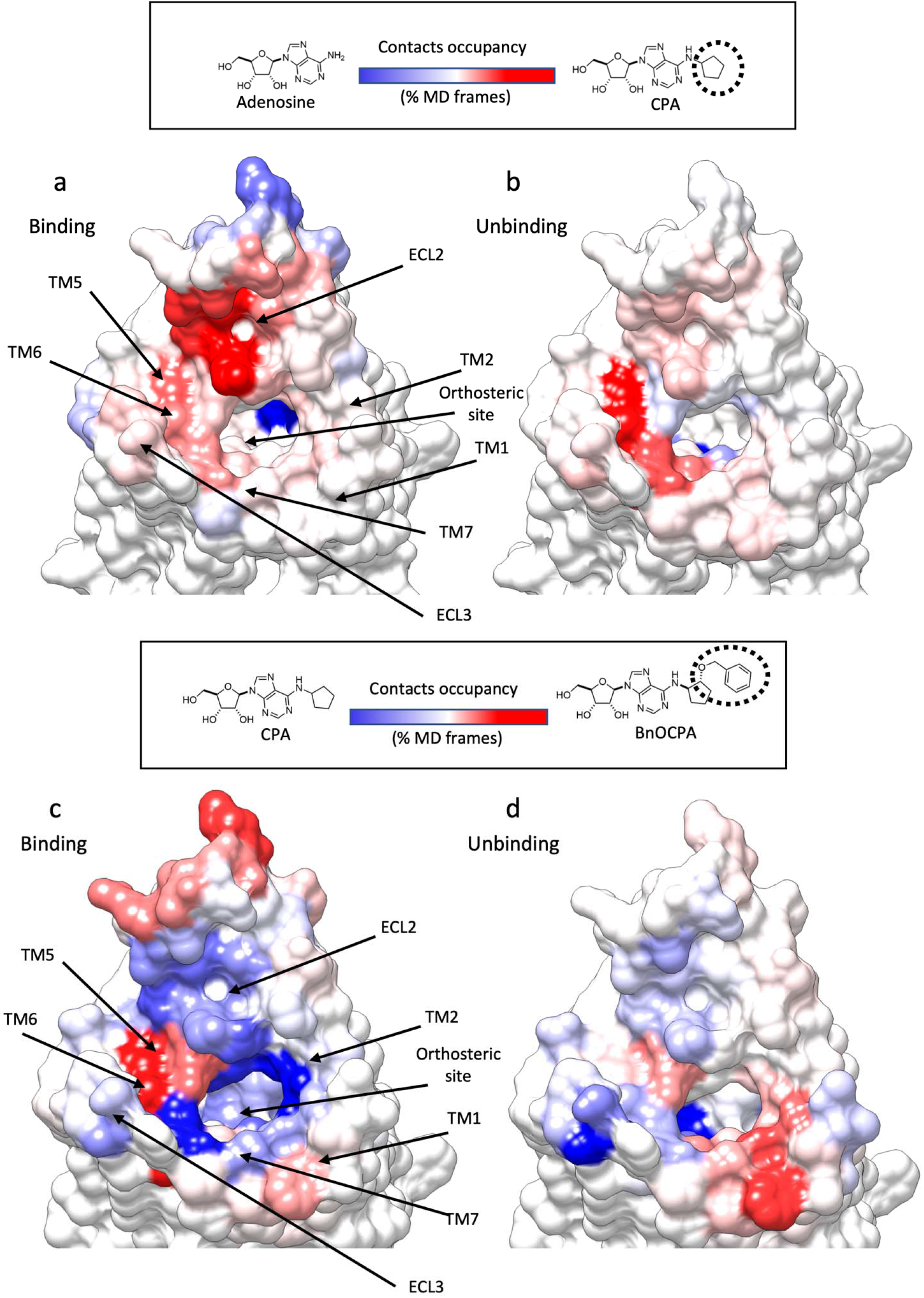
Structural modification of A_1_R agonists led to different simulated binding and unbinding mechanisms. **a)** and **b)** Heatmap showing the contact difference, during SuMD path sampling, between adenosine and CPA, which differ for the N6-cyclipentyl group (highlighted as a dashed circle line); A_1_R blue surfaces indicates more contacts formed with adenosine, while red surfaces are indicative of more contacts with CPA. **a**) Binding simulations; **b**) Unbinding simulations. **c)** and **d)** Heatmap showing the contacts difference, during SuMD path sampling, between CPA and BnOCPA, which differ for the oxybenzyl group (highlighted as a dashed circle line); A_1_R blue surfaces indicate more contacts formed with CPA, while red surfaces are indicative of more contacts with BnOCPA **c**) Binding simulations; **d**) Unbinding simulations.

#### 3.4.1 Introduction of the N6-cycloalkyl ring on the adenosine scaffold

In general, CPA formed more intermediate interactions than adenosine with the extracellular vestibule of A_1_R (Figure 4a,b). The presence of the N6-cycloalkyl substituent produced mode contacts with ECL2 and residues located at the top of TM1, TM5, TM6, TM7, and ECL3 (Figure 4a,b). This scenario is in good agreement with mutagenesis experiments (Table 2) showing CPA affinity (but not adenosine’s affinity) was significantly affected by T257^6.58^A, L258^6.59^A, H264^ECL3^A, and S267^7.32^A, all located in the proximity of the extracellular vestibule.

#### 3.4.2 Introduction of the 2-oxybenzyl group on CPA N6-cycloalkyl ring

The presence of this lipophilic moiety (BnOCPA was the bulkier ligand considered) favored more interactions with the top of TM5 and TM6 during association (Figure 4c) and with the top of TM1 and TM2 during dissociation (Figure 4d). CPA, on the other hand, was more prone to interact with the ECL2 (Figure 4c,d). The changes in the barycenter of the contacts during unbinding due to the *2-*oxybenzyl group could explain the unique profile that BnOCPA showed on A_1_R mutants F8^1.31^A and N70^2.65^A (Table 2). BnOCPA, indeed, was the only agonist significantly affected by N70^2.65^A, and the only one not affected by F8^1.31^A.

## 4. Conclusion

In the present work, unbiased nonequilibrium MD simulations and mutagenesis experiments were combined to study the dynamic binding and unbinding of A_1_R agonists. The *in silico* analysis suggested several involved A_1_R residues on regions so far poorly investigated^65^. Mutagenesis experiments generally confirmed the computational prediction and allow mapping novel receptor spots involved in the association or dissociation of the selective agonists.

The importance of intermediate metastable states along the (un)binding paths in modulating the overall affinity of ARs agonist is gradually emerging^12,66^. For the binding, our results indicated that A_1_R ECL2 is involved in numerous preliminary contacts, as well as the top of TM6, TM1, TM2 and TM7. The dissociation from A_1_R follows similar routes, but ECL2 is generally less engaged (especially by the more hydrophilic adenosine and NECA). The chemical modifications that increase the agonists’ selectivity towards A_1_R (e.g. the introduction of N6-cycloalkyl groups) also change the (un)binding mechanism, favoring the interactions with the extracellular vestibules in general, and the hydrophobic allosteric pocket located on ECL2 in particular.

GPCR-ligand complexes form and dissociate through multistep mechanisms. MD simulations and mutagenesis experiments are frequently combined to deliver structural insights on endpoint protein-ligand complexes. To the best of our knowledge^67–69^ this is the first that the whole process of formation/dissociation of several GPCR ligands is reconstructed with a combined *in silico* and *in vitro* approach. Our results pave the way to the rationalization of structure-kinetic relationship (SKR) for A_1_R, with potential repercussion on the rational design of long-awaited clinical agents.

## ASSOCIATED CONTENT

The following files are available free of charge:

Supporting Information (pdf)

Video S1-S4 (mp4)

## Author Contributions

The manuscript was written through contributions of all authors. All authors have given approval to the final version of the manuscript.

## Funding Sources

Leverhulme Trust (RPG-2017-255, CAR and GL to fund KB and GD).

## ACKNOWLEDGMENT

CAR is grateful for a Royal Society Industry Fellowship.

## Notes

### Competing Interest Statement

The authors have declared no competing interest.

