## Supporting Information (pdf) for "Deciphering the Agonist Binding Mechanism to the Adenosine A1 Receptor"

**Table of Contents**

Table S1-S5 pg 1-13
Figure S1-S5 pg 14-19
Captions Video S1-S4 pg 19

**Table S1. Contacts between $A_1R$ and the agonists during binding SuMD path sampling simulations.** Contact persistency is quantified as the percentage of frames (over all the frames obtained by merging the different replicas) in which protein residues were closer than 3.5 Å to the ligand. Residues in bold make direct contact with adenosine in the cryo-EM complex 6D9H.

| | | Occupancy (% MD frames) | | | | |
|---|---|---|---|---|---|---|
| | $A_1R$ Residue | Adenosine | NECA | CPA | HOCPA | BnOCPA |
| TM1 | Ser6 | 0.2 | 0.5 | 0.4 | 1 | 0.8 |
| | Ala7 | 0 | 0.1 | 0 | 0.1 | 0.1 |
| | Phe8 | 0.3 | 0.3 | 0.7 | 2.5 | 2.8 |
| | Gln9 | 2.1 | 3.3 | 5.2 | 2.9 | 3.1 |
| | Ala11 | 0 | 0 | 0 | 0.1 | 0.2 |
| | Tyr12 | 6.3 | 5.8 | 6.6 | 3.3 | 6 |
| | Ile13 | 0.1 | 0.4 | 0.8 | 0.2 | 0.6 |
| | Ile15 | 0.1 | 0 | 0 | 0.1 | 0.3 |
| | Glu16 | 2.9 | 1.9 | 2 | 1.7 | 2 |
| TM2 | Val58 | 0.1 | 0.4 | 1.2 | 0.3 | 0.4 |
| | Leu68 | 0 | 0.3 | 0 | 0 | 0 |
| | Val62 | 0.8 | 3.3 | 2.6 | 0.5 | 2.3 |
| | Ile63 | 0 | 0 | 0.8 | 0 | 0 |
| | Leu65 | 0.4 | 3.6 | 2.1 | 0.3 | 1.4 |
| | Ala66 | 4 | 5.4 | 6.9 | 3.7 | 3.1 |
| | Ile67 | 0 | 0 | 0.4 | 0.2 | 0 |
| | Ile69 | 6.1 | 9.8 | 9.7 | 7.5 | 2.8 |
| | Asn70 | 7 | 7.8 | 9.2 | 8.2 | 2 |
| | Ile71 | 0.3 | 0.9 | 1 | 0.7 | 0.1 |
| | Gly72 | 0.1 | 0.3 | 0.5 | 0.3 | 0 |
| ECL1 | Pro73 | 0.2 | 0.4 | 0.1 | 0.2 | 0.1 |
| | Gln74 | 1 | 2.8 | 2.1 | 1.5 | 2.7 |
| | Thr75 | 0.2 | 0.2 | 0 | 0 | 0.3 |
| | Tyr76 | 0.5 | 0.5 | 0 | 0 | 1.3 |
| | Phe77 | 0.3 | 0.5 | 1 | 0.2 | 0.6 |
| | His78 | 0 | 0.3 | 0.2 | 0 | 0.1 |
| | Thr79 | 0 | 0 | 0 | 0 | 0.1 |

| | | | | | | |
|---|---|---|---|---|---|---|
| TM3 | Cys80 | 0 | 0.4 | 0 | 0 | 0 |
| | Val83 | 0 | 0.6 | 0.1 | 0 | 0.2 |
| | Ala84 | 0.1 | 0.9 | 0.5 | 0.1 | 0.1 |
| | **Val87** | **9.5** | **11.4** | **8.6** | **9.3** | **6.5** |
| | Leu88 | 3 | 6.1 | 4.7 | 5.4 | 3.7 |
| | **Thr91** | **6.8** | **8.8** | **8.6** | **8.9** | **5.7** |
| | Gln92 | 0 | 0.7 | 0 | 0 | 0.1 |
| | Ile95 | 0 | 0.2 | 0 | 0.1 | 0 |
| TM4 | Met143 | 0 | 0.5 | 0.2 | 0 | 0 |
| | Phe144 | 0 | 0.3 | 0.4 | 0 | 0 |
| | Gly145 | 0 | 0.2 | 0.3 | 0.1 | 0.1 |
| ECL2 | Asn147 | 0.1 | 0.6 | 0.5 | 0.3 | 0.2 |
| | Asn148 | 4 | 6.7 | 15.2 | 11.5 | 10.8 |
| | Leu149 | 0 | 0.2 | 0.3 | 0 | 0.2 |
| | Ser150 | 0.3 | 0.4 | 0.4 | 0.6 | 0.6 |
| | Ala151 | 2.6 | 5 | 13.6 | 9.8 | 10.9 |
| | Val152 | 3.5 | 6.2 | 16.4 | 10.8 | 12.9 |
| | Glu153 | 0.4 | 0.2 | 0.1 | 0.3 | 0.6 |
| | Arg154 | 1.2 | 2.7 | 2.1 | 2.2 | 5 |
| | Ala155 | 3.4 | 6.3 | 13.2 | 10 | 11.6 |
| | Trp156 | 1.8 | 4.7 | 7.2 | 4.8 | 7.1 |
| | Ala157 | 0.7 | 0.8 | 0.3 | 0.5 | 2.7 |
| | Ala158 | 0.9 | 2.3 | 1.6 | 1.4 | 4.1 |
| | Asn159 | 2.2 | 5.3 | 8 | 5.6 | 7.8 |
| | Gly160 | 0.6 | 0.7 | 0.3 | 0.3 | 2.5 |
| | Ser161 | 1.1 | 2.5 | 1.4 | 0.7 | 4.1 |
| | Met162 | 1.2 | 1.4 | 0.4 | 0.4 | 4.8 |
| | Gly163 | 0.5 | 0.2 | 0 | 0.4 | 1.4 |
| | Glu164 | 0.2 | 0.1 | 0 | 0.1 | 0.6 |
| | Pro165 | 0.1 | 0.2 | 0.1 | 0.1 | 0.1 |
| | Val166 | 1.1 | 4.1 | 5 | 3.5 | 3.9 |
| | Ile167 | 2.2 | 3.8 | 8.7 | 5.5 | 5.7 |
| | Lys168 | 1.2 | 2 | 2.6 | 1.3 | 2 |

| | | | | | | |
|---|---|---|---|---|---|---|
| | Cys169 | 0 | 0.7 | 0.5 | 0 | 0.1 |
| | Glu170 | 1.9 | 3.5 | 5.8 | 6.6 | 1.2 |
| | **Phe171** | **21.1** | **18.7** | **22.9** | **19.3** | **17.8** |
| | **Glu172** | **15.7** | **19.1** | **20** | **23.2** | **24** |
| | Lys173 | 5.4 | 9.3 | 18.9 | 16.1 | 16.6 |
| | Val174 | 0.1 | 0.2 | 0.5 | 0 | 0.5 |
| | Ile175 | 2 | 3.6 | 5.4 | 4 | 10.3 |
| TM5 | Ser176 | 2.1 | 2.6 | 4 | 2 | 1.7 |
| | Met177 | 7 | 6.5 | 16 | 15.2 | 21.9 |
| | Glu178 | 2 | 1.4 | 2.4 | 2.1 | 1.2 |
| | **Met180** | **9.7** | **10.2** | **13.7** | **12.9** | **14.2** |
| | Val181 | 0.1 | 0 | 0.8 | 0.1 | 0.1 |
| | Tyr182 | 0.3 | 0.5 | 0.2 | 0.8 | 0.2 |
| | Asn184 | 0.8 | 3 | 3.6 | 2.8 | 1.5 |
| | Val189 | 0 | 0.2 | 0.5 | 0 | 0 |
| TM6 | Ser246 | 0.1 | 0 | 0.6 | 0.1 | 0.1 |
| | **Trp247** | 2.7 | 3 | 3.3 | 3.2 | 4.5 |
| | **Leu250** | 13.4 | 13.6 | 16.4 | 17.2 | 16.2 |
| | His251 | 3.6 | 4 | 8 | 4.8 | 3.7 |
| | Leu253 | 0.6 | 0.6 | 2.1 | 5.4 | 7.2 |
| | **Asn254** | 11.2 | 11 | 15.1 | 18.1 | 21.4 |
| | Ile256 | 0 | 0.3 | 0 | 0.2 | 0.1 |
| | Thr257 | 3.8 | 3.2 | 10.5 | 12.2 | 16.1 |
| | Leu258 | 2.4 | 2 | 4.4 | 3.8 | 4.8 |
| | Phe259 | 0.7 | 1 | 0.3 | 1.2 | 0.5 |
| | Cys260 | 1.1 | 0.4 | 1.1 | 0.4 | 0.4 |
| ECL3 | Pro261 | 1.8 | 3.2 | 4.9 | 4.9 | 4.6 |
| | Ser262 | 1.4 | 1.8 | 3.9 | 2.3 | 2.3 |
| | Cys263 | 2.2 | 3.4 | 4.4 | 3.9 | 1.3 |
| | His264 | 2.1 | 4.2 | 6.2 | 7.9 | 1.7 |
| | Lys265 | 6.6 | 6.8 | 12.9 | 12.8 | 6.3 |
| | Pro266 | 0.4 | 0.3 | 0.2 | 1 | 0.7 |
| | Ser267 | 3.2 | 4.9 | 5 | 8.6 | 2.2 |

| | | | | | | |
|---|---|---|---|---|---|---|
| TM7 | Ile268 | 0 | 0 | 0.1 | 1.1 | 0.8 |
| | Thr270 | 4.8 | 8.3 | 8.9 | 12 | 10.5 |
| | Tyr271 | 5 | 8 | 7.8 | 10.3 | 5.3 |
| | Ile272 | 0 | 0 | 0 | 0 | 0.2 |
| | Ala273 | 0 | 0 | 1.6 | 0 | 0 |
| | **Ile274** | **11.7** | **16.1** | **13.3** | **10.9** | **9.1** |
| | Phe275 | 0 | 0 | 0 | 0 | 0.4 |
| | **Ser277** | **4.6** | **5** | **5.4** | **2.4** | **4.1** |
| | **His278** | **7.4** | **4.9** | **4.8** | **4.1** | **4.9** |

**Table S2. Contacts between A$_1$R and the agonists during unbinding SuMD path sampling simulations.** Contact persistency is quantified as the percentage of frames (over all the frames obtained by merging the different replicas) in which protein residues were closer than 3.5 Å to the ligand. Residues in bold make direct contact with adenosine in the cryo-EM complex 6D9H.

| | | Occupancy (% MD frames) | | | | |
|---|---|---|---|---|---|---|
| | Residue | Adenosine | NECA | CPA | HOCPA | BnOCPA |
| TM1 | Ser6 | 2.1 | 2.6 | 2.8 | 2.6 | 3.9 |
| | Ala7 | 0.6 | 0.3 | 1.1 | 0.2 | 1.2 |
| | Phe8 | 1 | 2.7 | 5.2 | 3.2 | 14.5 |
| | Gln9 | 3.6 | 9.9 | 7 | 5.9 | 12.9 |
| | Ala10 | 0 | 0.8 | 0.8 | 0.1 | 0.1 |
| | Ala11 | 0 | 0.1 | 0.2 | 0 | 1.1 |
| | Tyr12 | 5.1 | 13.1 | 5.9 | 7 | 17.8 |
| | Ile13 | 0 | 4.2 | 0.7 | 0.9 | 1.2 |
| | Ile15 | 0 | 0.2 | 0.2 | 0 | 1.1 |
| | Glu16 | 1.4 | 6.5 | 2.7 | 2.3 | 3.8 |
| TM2 | Val58 | 0.2 | 1 | 1.5 | 0.1 | 0.8 |
| | Gly59 | 0 | 0 | 0 | 0 | 0 |
| | Leu61 | 0 | 0.3 | 0 | 0 | 0 |
| | Val62 | 1.4 | 5.6 | 4 | 0 | 1.2 |
| | Ile63 | 0.1 | 0.1 | 0.6 | 0 | 1 |
| | Pro64 | 0 | 0 | 0 | 0 | 0.1 |
| | Leu65 | 0.1 | 5 | 0.4 | 0 | 0.6 |
| | Ala66 | 9.5 | 9.9 | 11.5 | 5.7 | 8.3 |
| | Ile67 | 0 | 0.1 | 0.2 | 0.5 | 1.9 |
| | Leu68 | 0 | 0 | 0 | 0 | 0.2 |
| | Ile69 | 11.1 | 20.4 | 8.2 | 6.7 | 8.1 |
| | Asn70 | 11.5 | 14.2 | 14.6 | 11.4 | 14.8 |
| | Ile71 | 1.4 | 0.9 | 1.2 | 1.7 | 1.3 |
| | Gly72 | 0.5 | 1.8 | 0.1 | 0.3 | 0.2 |
| ECL1 | Pro73 | 0.1 | 1 | 0 | 0.1 | 0.2 |
| | Gln74 | 0.7 | 0.8 | 1.1 | 0.2 | 2.1 |
| | Thr75 | 0 | 0 | 0 | 0 | 0.3 |

| | | | | | | |
|---|---|---|---|---|---|---|
| TM3 | Tyr76 | 0.1 | 0.1 | 0.1 | 0 | 0.7 |
| | Phe77 | 0 | 0.5 | 0.7 | 0.2 | 0.3 |
| | Val83 | 0 | 0.3 | 0 | 0 | 0.3 |
| | Ala84 | 0 | 1.3 | 0 | 0.4 | 0.6 |
| | **Val87** | **18.6** | **28.5** | **19.6** | **17.2** | **13.4** |
| | Leu88 | 2.5 | 14.2 | 4.4 | 8.7 | 4.9 |
| | **Thr91** | **17.3** | **28.6** | **18** | **17.6** | **8.6** |
| | Gln92 | 0 | 13.7 | 0.2 | 0.7 | 0.1 |
| | Ile95 | 0 | 6.7 | 0 | 0.1 | 0.1 |
| TM4 | Val138 | 0 | 0.2 | 0 | 0 | 0 |
| | Pro142 | 0 | 0.2 | 0 | 0 | 0 |
| | Gly145 | 0.2 | 0 | 0 | 0 | 0 |
| ECL2 | Asn147 | 0.2 | 0.5 | 0.2 | 0 | 0.1 |
| | Asn148 | 0.3 | 0.5 | 4.4 | 1.5 | 2.3 |
| | Leu149 | 0.1 | 0.2 | 0.1 | 0 | 0 |
| | Ser150 | 0.1 | 0.1 | 0.2 | 0 | 0.1 |
| | Ala151 | 0.2 | 0.7 | 4.4 | 1.5 | 2.6 |
| | Val152 | 0.4 | 1.2 | 6.5 | 1.7 | 3.9 |
| | Glu153 | 0.1 | 0.4 | 0.1 | 0 | 0.2 |
| | Arg154 | 0.3 | 0.5 | 1 | 0.4 | 0.9 |
| | Ala155 | 0.5 | 1.1 | 6.6 | 1.9 | 3.4 |
| | Trp156 | 0.7 | 0.8 | 4.4 | 1.1 | 3.5 |
| | Ala157 | 0.2 | 0.4 | 0.1 | 0 | 0.4 |
| | Ala158 | 0.3 | 0.6 | 0.7 | 0.4 | 0.7 |
| | Asn159 | 0.6 | 0.8 | 4.5 | 1.3 | 2.7 |
| | Gly160 | 0.2 | 0.1 | 0.2 | 0 | 0.4 |
| | Ser161 | 0.4 | 0.2 | 0.7 | 0.2 | 1.9 |
| | Met162 | 0.6 | 0.2 | 0.2 | 0 | 2.1 |
| | Gly163 | 0.1 | 0.3 | 0.1 | 0 | 0.3 |
| | Glu164 | 0.1 | 0.2 | 0 | 0 | 0.1 |
| | Pro165 | 0 | 0.3 | 0.1 | 0 | 0.1 |
| | Val166 | 0.6 | 0.7 | 2.9 | 0.4 | 2.8 |
| | Ile167 | 0.4 | 0.7 | 3.6 | 0.8 | 2.3 |

| | | | | | | |
|---|---|---|---|---|---|---|
| | Lys168 | 1.7 | 2.3 | 1.6 | 1.9 | 1.6 |
| | Cys169 | 0 | 2.2 | 0 | 0 | 0.1 |
| | Glu170 | 3.8 | 4.6 | 3 | 2.7 | 2.5 |
| | **Phe171** | **37.2** | **50.4** | **38.2** | **44.1** | **26.9** |
| | **Glu172** | **23.1** | **32.9** | **21.3** | **33.7** | **20.7** |
| | Lys173 | 0.8 | 1.6 | 7.2 | 3 | 4 |
| | Val174 | 0 | 0.5 | 0.3 | 0.1 | 0 |
| | Ile175 | 0 | 7.8 | 7.1 | 4.2 | 12.5 |
| TM5 | Ser176 | 0.1 | 1 | 1.5 | 0.8 | 1.8 |
| | Met177 | 2.1 | 10.5 | 27.2 | 33.3 | 22.4 |
| | Glu178 | 0.1 | 0.3 | 1.3 | 3.4 | 0.2 |
| | TYr179 | 0 | 0.4 | 0 | 0 | 0 |
| | **Met180** | **17** | **30.9** | **16.2** | **22.9** | **18.1** |
| | Val181 | 0 | 0.4 | 2 | 4.8 | 0.2 |
| | Tyr182 | 0 | 0 | 0.1 | 3.7 | 0.1 |
| | Asn184 | 0.8 | 12.6 | 0.9 | 1 | 1.2 |
| | Phe185 | 0 | 0 | 0 | 0.4 | 0 |
| | Phe186 | 0 | 0 | 0 | 0.5 | 0 |
| | Val189 | 0 | 2.8 | 0 | 0 | 0 |
| TM6 | Phe243 | 0 | 0.3 | 0 | 0 | 0 |
| | **Trp247** | **7.1** | **13.9** | **0** | **0.1** | **5.2** |
| | **Leu250** | **24.4** | **35.9** | **27.8** | **35.6** | **17.1** |
| | His251 | 3.7 | 11.1 | 1.8 | 2.3 | 3 |
| | Leu253 | 0 | 2.8 | 13.8 | 20.5 | 9.5 |
| | **Asn254** | **25.8** | **33.6** | **34.7** | **47.4** | **22.7** |
| | Cys255 | 0 | 0 | 0 | 0.3 | 0 |
| | ILe256 | 0 | 0 | 1.7 | 3.2 | 0 |
| | Thr257 | 0.1 | 3 | 24.2 | 33.5 | 15.7 |
| | Leu258 | 0.1 | 1.1 | 3.4 | 7.2 | 3.4 |
| | Phe259 | 0 | 0.3 | 0.3 | 1.8 | 0.1 |
| | Cys260 | 0 | 0.3 | 1.6 | 3 | 0 |
| | Pro261 | 0.3 | 0.7 | 1.2 | 1.5 | 0.3 |
| | Ser262 | 0.5 | 1 | 3.6 | 4.9 | 0.7 |

| | | | | | | |
|---|---|---|---|---|---|---|
| ECL3 | Cys263 | 0.5 | 1.2 | 6.5 | 11.3 | 0.5 |
| | His264 | 2.1 | 1.5 | 4.7 | 7.6 | 2.9 |
| | Lys265 | 5.3 | 5.2 | 22.8 | 39.9 | 14.7 |
| TM7 | Pro266 | 0.3 | 0.6 | 2.8 | 6.2 | 3 |
| | Ser267 | 5.2 | 3.3 | 7.9 | 5.6 | 10.8 |
| | Ile268 | 0.1 | 0.2 | 1.6 | 0.5 | 4.3 |
| | Leu269 | 0 | 0 | 0.2 | 0 | 0.1 |
| | Thr270 | 7.8 | 8.7 | 22.2 | 29.2 | 22 |
| | Tyr271 | 7.2 | 11.9 | 13.1 | 11 | 19.3 |
| | Ile272 | 0 | 0 | 0.1 | 0 | 0.5 |
| | **Ile274** | **26.5** | **30.8** | **20.8** | **18.8** | **17.8** |
| | Phe275 | 0 | 0.2 | 0.1 | 0 | 0.7 |
| | **Ser277** | **17.9** | **17.1** | **3.1** | **1.9** | **4.7** |
| | **His278** | **12.7** | **17** | **11.2** | **6.6** | **3** |

**Table S3. Hydrogen bonds between $A_1R$ and the agonists during binding SuMD path sampling simulations.** Hydrogen bond persistency is quantified as the percentage of frames (over all the frames obtained by merging the different replicas) in which protein residues were closer than 3.5 Å to the ligand. Residues in bold make hydrogen bond with adenosine in the cryo-EM complex 6D9H.

| | Occupancy (% MD frames) | | | | | |
|---|---|---|---|---|---|---|
| | Residue | Adenosine | NECA | CPA | HOCPA | BnOCPA |
| TM1 | Ile5 | 0 | 0.1 | 0 | 0.1 | 0 |
| | Ser6 | 0.1 | 0.1 | 0 | 0.2 | 0.2 |
| | Gln9 | 1.2 | 1.1 | 2.4 | 1.1 | 0.8 |
| | Tyr12 | 1.7 | 2.6 | 0.7 | 0.7 | 0.7 |
| | Glu16 | 2.2 | 1.7 | 1.2 | 1.3 | 1.6 |
| TM2 | Asn70 | 3.8 | 4.1 | 3.7 | 2.1 | 0.4 |
| ECL1 | Gln74 | 0.3 | 1 | 0.9 | 0.6 | 0.7 |
| | Thr75 | 0.1 | 0.1 | 0 | 0 | 0 |
| TM3 | Tyr76 | 0.1 | 0.1 | 0 | 0 | 0.2 |
| | His78 | 0 | 0.1 | 0 | 0 | 0 |
| | Thr91 | 2 | 4.3 | 4.1 | 3.6 | 2 |
| ECL2 | Asn147 | 0 | 0.3 | 0.2 | 0 | 0 |
| | Asn148 | 1.5 | 2.8 | 2.4 | 2.6 | 1.7 |
| | Ser150 | 0.1 | 0.1 | 0.1 | 0.1 | 0.1 |
| | Glu153 | 0.3 | 0.1 | 0 | 0.2 | 0.2 |
| | Arg154 | 0.2 | 0.6 | 0.3 | 0.4 | 0.6 |
| | Asn159 | 1 | 2.5 | 2.8 | 2 | 1.9 |
| | Ser161 | 0.2 | 1.1 | 0.5 | 0.2 | 0.8 |
| | Glu164 | 0.1 | 0 | 0 | 0 | 0.1 |
| | Lys168 | 0.1 | 0.3 | 0.7 | 0.5 | 0.1 |
| | Glu170 | 1.4 | 1.5 | 2.5 | 2.3 | 0.3 |
| | **Glu172** | **8.6** | **10** | **5.1** | **8.1** | **4.1** |
| | Lys173 | 0.5 | 0.7 | 1.3 | 2 | 1 |
| TM5 | Ser176 | 0.9 | 0.7 | 1.5 | 0.4 | 0.2 |
| | Glu178 | 1.6 | 1 | 1.5 | 0.7 | 0.6 |
| | Tyr182 | 0.1 | 0.2 | 0 | 0.2 | 0 |
| | Asn184 | 0.5 | 1 | 2.4 | 2.4 | 0.5 |

| | | | | | | |
|---|---|---|---|---|---|---|
| TM6 | Trp247 | 0 | 0 | 0.3 | 0 | 0 |
| | His251 | 1.2 | 0.3 | 3.9 | 0.4 | 0.4 |
| | **Asn254** | **9.4** | **8.4** | **9.2** | **14.3** | **16.5** |
| | Thr257 | 0.2 | 0.2 | 0.9 | 1.6 | 1.4 |
| ECL3 | Ser262 | 0.1 | 0.2 | 0.3 | 0.3 | 0.3 |
| | His264 | 0.5 | 0.9 | 1 | 2.1 | 0.3 |
| | Lys265 | 1.4 | 1.2 | 1.9 | 2.5 | 1.2 |
| TM7 | Ser267 | 1 | 1.9 | 0.7 | 2.4 | 0.6 |
| | The270 | 0.7 | 0.5 | 0.6 | 1.2 | 0.6 |
| | Tyr271 | 1.1 | 1.9 | 0.7 | 1.3 | 0.5 |
| | Ser277 | 2 | 1 | 1.4 | 0.4 | 0.9 |
| | His278 | 0.5 | 0.3 | 0.6 | 0.2 | 0.5 |

**Table S4. Hydrogen bonds between $A_1R$ and the agonists during binding SuMD path sampling simulations.** Hydrogen bond persistency is quantified as the percentage of frames (over all the frames obtained by merging the different replicas) in which protein residues were closer than 3.5 Å to the ligand. Residues in bold make hydrogen bond with adenosine in the cryo-EM complex 6D9H

| | | Occupancy (% MD frames) | | | | |
|---|---|---|---|---|---|---|
| | Residue | Adenosine | NECA | CPA | HOCPA | BnOCPA |
| TM1 | Ile5 | 0.2 | 0.6 | 0.1 | 0.2 | 0.2 |
| | Ser6 | 0.2 | 0.2 | 0.5 | 0.4 | 0.9 |
| | Gln9 | 0.6 | 3.2 | 2.9 | 2 | 5.8 |
| | Tyr12 | 0 | 0.9 | 1.9 | 2.3 | 3.6 |
| | Glu16 | 0 | 0.8 | 2 | 1.2 | 3.4 |
| TM2 | Asn70 | 0.4 | 4.1 | 5.1 | 3.6 | 5.9 |
| ECL1 | Gln74 | 0.2 | 0.1 | 0.3 | 0.1 | 0.5 |
| | Thr75 | 0 | 0 | 0 | 0 | 0.2 |
| TM3 | Tyr76 | 0 | 0 | 0 | 0 | 0.1 |
| | Thr91 | 0.1 | 23.1 | 6.2 | 5.5 | 0.8 |
| | Gln92 | 0 | 0.1 | 0.1 | 0.3 | 0 |
| | Asn147 | 0.1 | 0 | 0 | 0 | 0 |

| | | | | | | |
|---|---|---|---|---|---|---|
| ECL2 | Asn148 | 0 | 0.1 | 1.1 | 0.3 | 0.8 |
| | Arg154 | 0 | 0 | 0.2 | 0 | 0.1 |
| | Asn159 | 0.1 | 0.1 | 1.6 | 0.6 | 0.6 |
| | Ser161 | 0 | 0 | 0.2 | 0 | 0.2 |
| | Lys168 | 0.2 | 0.2 | 0.7 | 0.8 | 0.3 |
| | Glu164 | 0.1 | 0.8 | 1.8 | 1.5 | 1.8 |
| | **Glu172** | **15.9** | **13.4** | **1.8** | **11.9** | **0.5** |
| | Lys173 | 0.1 | 0.1 | 0.3 | 0.7 | 0.2 |
| TM5 | Ser176 | 0 | 0 | 0 | 0.1 | 0 |
| | Glu178 | 0 | 0.1 | 0.3 | 0.5 | 0.1 |
| | Tyr182 | 0 | 0 | 0 | 0.8 | 0.1 |
| | Asn184 | 0 | 0.4 | 0.7 | 0.7 | 0.9 |
| TM6 | His251 | 0.2 | 0.1 | 0.1 | 0.3 | 0.3 |
| | **Asn254** | **24.8** | **23.4** | **20.2** | **35.5** | **15.4** |
| | The257 | 0 | 0.5 | 2.3 | 7.4 | 0.1 |
| ECL3 | Ser262 | 0 | 0.1 | 0.5 | 0.6 | 0.1 |
| | His264 | 0.3 | 0.1 | 0.7 | 0.3 | 0.3 |
| | Lys265 | 0.5 | 0.5 | 2.4 | 21.1 | 3.2 |
| TM7 | Ser267 | 1.3 | 1.3 | 2.3 | 0.9 | 1.1 |
| | Thr270 | 0.2 | 0.9 | 0.2 | 1.7 | 1.7 |
| | Tyr271 | 0.3 | 1.3 | 1.9 | 2.3 | 3.6 |
| | Ser277 | 0 | 0 | 0.6 | 0.6 | 2.4 |
| | His278 | 0 | 0.4 | 0.9 | 0.7 | 1.1 |

**Table S5. NanoBRET competition binding essays in WT and mutant Nluc-A1R.**

NECA affinity (pKi) at WT and mutant Nluc-$A_1R$.

| | **$pK_i$** | |
|---|---|---|
| | **NECA** | ***n*** |
| **WT** | 6.39 ±0.04 | 4 |
| **I175A** | 5.51 ±0.07**** | 2 |
| **T257A** | 5.91 ±0.04**** | 4 |
| **L258A** | 5.66 ±0.07**** | 5 |
| **S267A** | 6.09 ±0.03* | 2 |
| **Y271A** | 5.98 ±0.05*** | 3 |
| **L253T** | 5.92 ±0.11** | 3 |
| **L253F** | 5.85 ±0.02**** | 3 |
| **L253G** | - | |

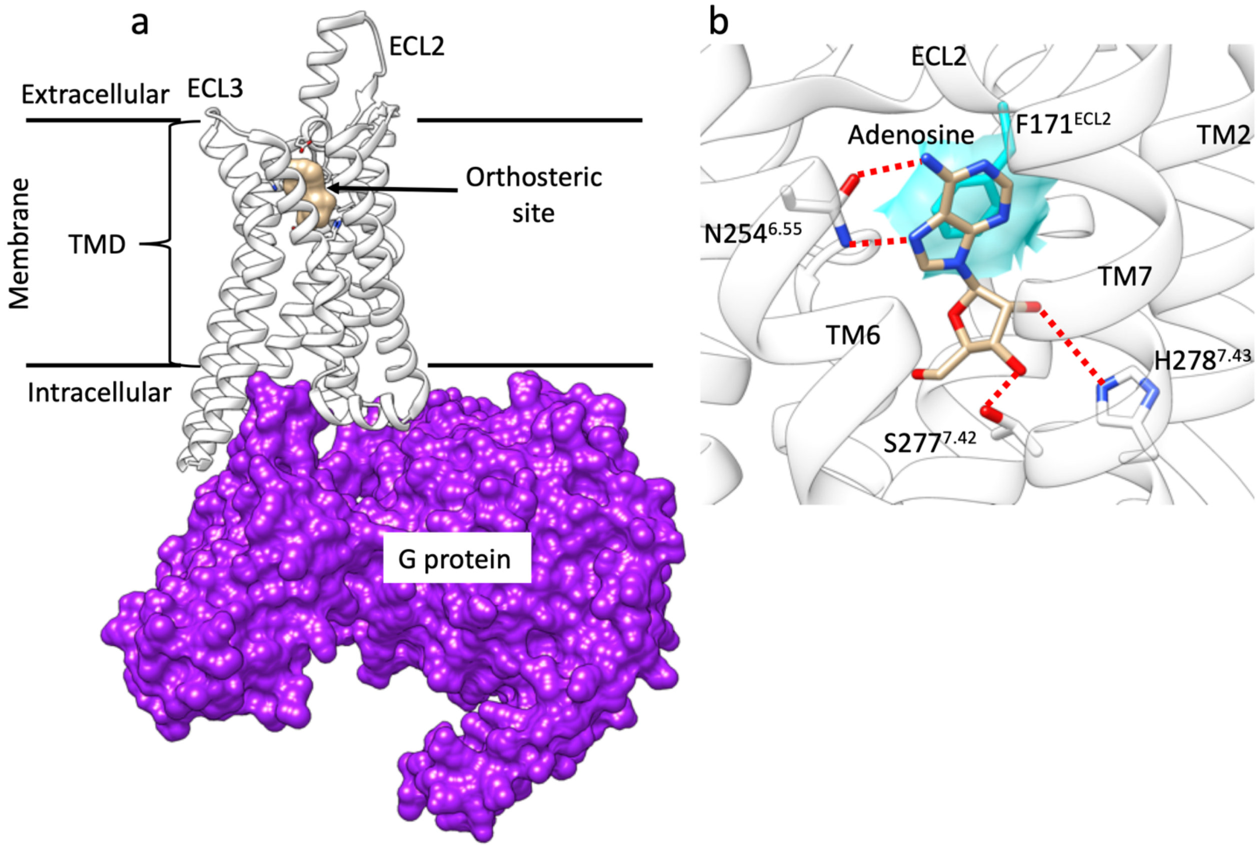


Figure S1. a) $A_1AR$ (white ribbon) transmembrane domain (TMD) spans the cytosolic membrane seven times and shaped the orthosteric binding site and the G protein binding site (purple surface); b) magnification of the adenosine (tan stick representation) bound to the orthosteric site of $A_1AR$. Hydrogen bonds are shown as dashed lines, while hydrophobic contacts are depicted as cyan transparent surfaces.

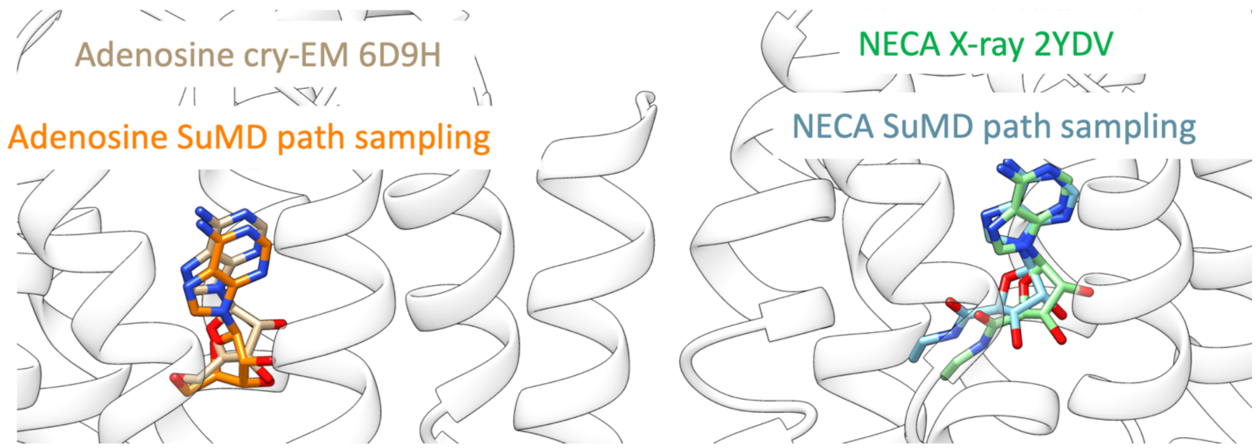

Figure S2. Comparison between orthosteric binding modes on $A_1R$ (white ribbon) sampled by means of SuMD path sampling and the experimental conformations. Adenosine is shown on left hand panel, while and NDECA on the right panel.

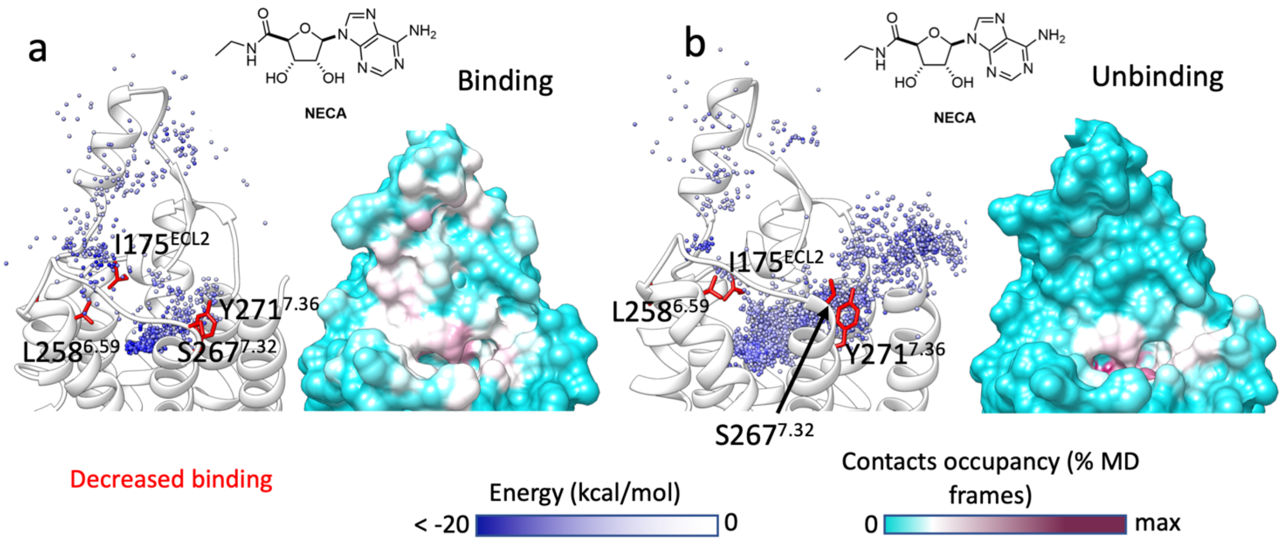


Figure S3. Simulated binding and unbinding paths of NECA and the relative position of the $A_1R$ mutants tested. a) Left hand panels, position of NECA centroid during binding simulations, colored according to the interaction energy with $A_1R$ (white ribbon); residues mutated are shown as stick and colored according to effect on the affinity (red: decreased affinity). Right hand panels, $A_1R$-agonist contacts plotted onto the protein surface and colored according to the contacts occupancy. b) Left hand panels, position of NECA centroid during unbinding simulations, colored according to the interaction energy with $A_1R$ (white ribbon); residues mutated are shown as stick and colored according to effect on the affinity (red: decreased affinity). Right hand panels, $A_1R$-agonist contacts plotted onto the protein surface and colored according to the contacts occupancy

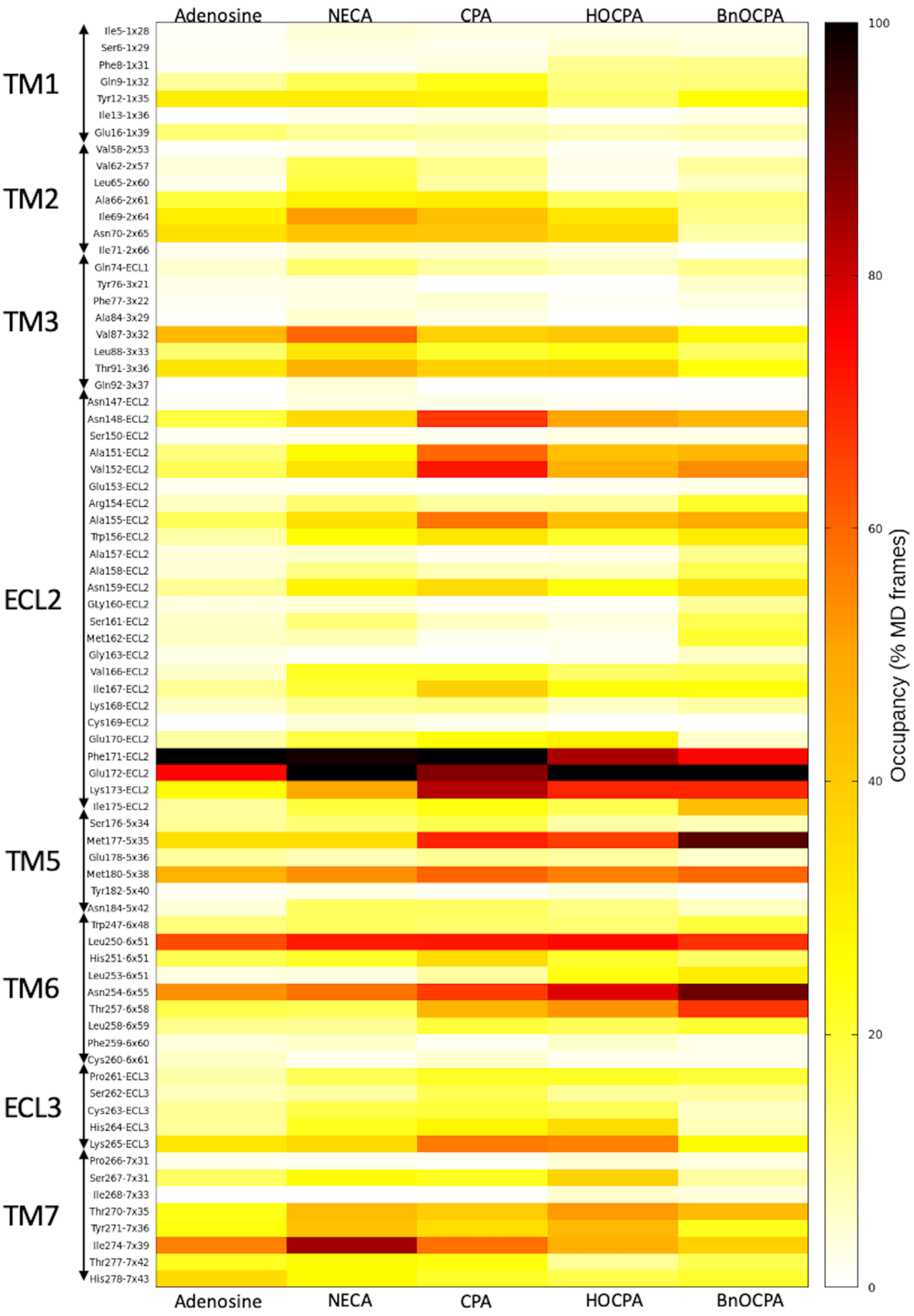
Adenosine
NECA
CPA
HOCPA
BnOCPA
TM1
TM2
TM3
ECL2
TM5
TM6
ECL3
TM7
Ile5-1x28
Ser6-1x29
Phe8-1x31
Gln9-1x32
Tyr12-1x35
Ile13-1x36
Glu16-1x39
Val58-2x53
Val62-2x57
Leu65-2x60
Ala66-2x61
Ile69-2x64
Asn70-2x65
Ile71-2x66
Gln74-ECL1
Tyr76-3x21
Phe77-3x22
Ala84-3x29
Val87-3x32
Leu88-3x33
Thr91-3x36
Gln92-3x37
Asn147-ECL2
Asn148-ECL2
Ser150-ECL2
Ala151-ECL2
Val152-ECL2
Glu153-ECL2
Arg154-ECL2
Ala155-ECL2
Trp156-ECL2
Ala157-ECL2
Ala158-ECL2
Asn159-ECL2
Gly160-ECL2
Ser161-ECL2
Met162-ECL2
Gly163-ECL2
Val166-ECL2
Ile167-ECL2
Lys168-ECL2
Cys169-ECL2
Glu170-ECL2
Phe171-ECL2
Glu172-ECL2
Lys173-ECL2
Ile175-ECL2
Ser176-5x34
Met177-5x35
Glu178-5x36
Met180-5x38
Tyr182-5x40
Asn184-5x42
Trp247-6x48
Leu250-6x51
His251-6x51
Leu253-6x51
Asn254-6x55
Thr257-6x58
Leu258-6x59
Phe259-6x60
Cys260-6x61
Pro261-ECL3
Ser262-ECL3
Cys263-ECL3
His264-ECL3
Lys265-ECL3
Pro266-7x31
Ser267-7x31
Ile268-7x33
Thr270-7x35
Tyr271-7x36
Ile274-7x39
Thr277-7x42
His278-7x43
Adenosine
NECA
CPA
HOCPA
BnOCPA
100
80
60
40
20
0
Occupancy (% MD frames)

Figure S4. Heatmap showing the $A_1R$ contacts with adenosine, NECA, CPA, HOCPA, and BnOCPA during binding SuMD path sampling. For each agonist, data are reported as the percentage of MD frames (occupancy) normalized for the maximum occupancy occurred.

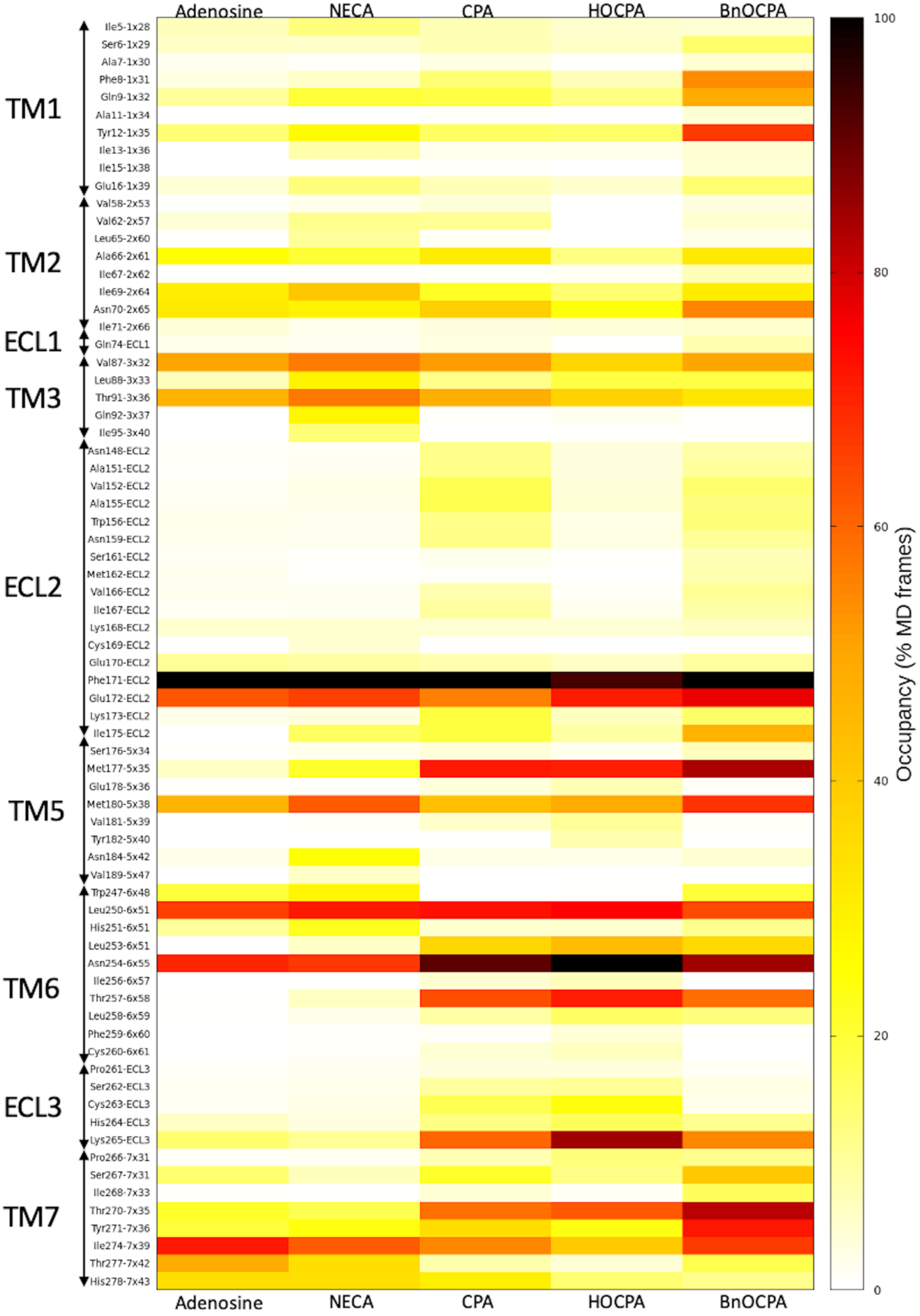
Adenosine
NECA
CPA
HOCPA
BnOCPA
TM1
TM2
ECL1
TM3
ECL2
TM5
TM6
ECL3
TM7
Ile5-1x28
Ser6-1x29
Ala7-1x30
Phe8-1x31
Gln9-1x32
Ala11-1x34
Tyr12-1x35
Ile13-1x36
Ile15-1x38
Glu16-1x39
Val58-2x53
Val62-2x57
Leu65-2x60
Ala66-2x61
Ile67-2x62
Ile69-2x64
Asn70-2x65
Ile71-2x66
Gln74-ECL1
Val87-3x32
Leu88-3x33
Thr91-3x36
Gln92-3x37
Ile95-3x40
Asn148-ECL2
Ala151-ECL2
Val152-ECL2
Ala155-ECL2
Trp156-ECL2
Asn159-ECL2
Ser161-ECL2
Met162-ECL2
Val166-ECL2
Ile167-ECL2
Lys168-ECL2
Cys169-ECL2
Glu170-ECL2
Phe171-ECL2
Glu172-ECL2
Lys173-ECL2
Ile175-ECL2
Ser176-5x34
Met177-5x35
Glu178-5x36
Met180-5x38
Val181-5x39
Tyr182-5x40
Asn184-5x42
Val189-5x47
Trp247-6x48
Leu250-6x51
His251-6x51
Leu253-6x51
Asn254-6x55
Ile256-6x57
Thr257-6x58
Leu258-6x59
Phe259-6x60
Cys260-6x61
Pro261-ECL3
Ser262-ECL3
Cys263-ECL3
His264-ECL3
Lys265-ECL3
Pro266-7x31
Ser267-7x31
Ile268-7x33
Thr270-7x35
Tyr271-7x36
Ile274-7x39
Thr277-7x42
His278-7x43
Adenosine
NECA
CPA
HOCPA
BnOCPA
100
80
60
40
20
0
Occupancy (% MD frames)

Figure S5. Heatmap showing the $A_1R$ contacts with adenosine, NECA, CPA, HOCPA, and BnOCPA during unbinding SuMD path sampling. For each agonist, data are reported as the percentage of MD frames (occupancy) normalized for the maximum occupancy occurred.

**Caption Video S1**

Merged SuMD simulations of adenosine binding (left side) and unbinding (right side). $A_1R$ is represented as white ribbon, while adenosine as van der Waals spheres. $A_1R$ residues located within 4 Å from the ligand are shown as stick. The time pace showed in the binding and unbinding simulations is different (unbinding simulation time is faster).

**Caption Video S2**

Merged SuMD simulations of NECA binding (left side) and unbinding (right side). $A_1R$ is represented as white ribbon, while NECA as van der Waals spheres. $A_1R$ residues located within 4 Å from the ligand are shown as stick. The time pace showed in the binding and unbinding simulations is different (unbinding simulation time is faster).

**Caption Video S3**

Merged SuMD simulations of CPA binding (left side) and unbinding (right side). $A_1R$ is represented as white ribbon, while CPA as van der Waals spheres. $A_1R$ residues located within 4 Å from the ligand are shown as stick. The time pace showed in the binding and unbinding simulations is different (unbinding simulation time is faster).

**Caption Video S4**

Merged SuMD simulations of BnOCPA binding (left side) and unbinding (right side). $A_1R$ is represented as white ribbon, while BnOCPA as van der Waals spheres. $A_1R$ residues located within 4 Å from the ligand are shown as stick. The time pace showed in the binding and unbinding simulations is different (unbinding simulation time is faster).
